# Sleep cycle-dependent vascular dynamics enhance perivascular cerebrospinal fluid flow and solute transport

**DOI:** 10.1101/2022.07.14.500017

**Authors:** Laura Bojarskaite, Daniel M. Bjørnstad, Alexandra Vallet, Kristin M. Gullestad Binder, Céline Cunen, Kjell Heuser, Miroslav Kuchta, Kent-Andre Mardal, Rune Enger

**Affiliations:** GliaLab and the Letten Centre, Division of Anatomy, Department of Molecular Medicine, Institute of Basic Medical Sciences, University of Oslo, 0317 Oslo, Norway; Department of Neurology, Oslo University Hospital, 0027 Oslo, Norway; Department of Mathematics, University of Oslo, 0316, Oslo, Norway; Norwegian Computing Center, 0314 Oslo, Norway; Department of Numerical Analysis and Scientific Computing, Simula Research Laboratory, Kristian Augusts gate 23, 0134 Oslo, Norway

**Author notes:** These authors contributed equally to this work (Laura Bojarskaite, Daniel M. Bjørnstad, Alexandra Vallet).

**Keywords:** glymphatic, endfoot, astrocyte, pial artery, penetrating arteriole

## Abstract

Perivascular spaces (PVS) are important highways for fluid and solute transport in the brain enabling efficient waste clearance during sleep. Using two-photon imaging of naturally sleeping mice we demonstrate sleep cycle-dependent PVS dynamics – slow, large-amplitude oscillations in NREM, a reduction in REM and an enlargement upon awakening at the end of a sleep cycle. By biomechanical modeling we demonstrate that these sleep cycle-dependent PVS dynamics drive fluid flow and solute transport.

## MAIN

Perivascular spaces (PVS) lined by the astrocytic endfeet, are key passageways for movement and exchange of fluids and solutes, and play important roles for drug delivery into the brain and waste clearance^1,2^. A current model for brain waste clearance – the glymphatic system – states that cerebrospinal fluid (CSF) flows along pial arteries, enters the brain via PVS of penetrating arterioles, then flows through the parenchyma collecting extracellular waste, before it exits in PVS along veins or arteries^3^. This process is thought to be facilitated by mechanical forces created by the vasculature^4^, such as heartbeat driven pial artery pulsations seen in experiments with anesthetized mice^5^ or vasomotion of longer time scales observed in wakefulness^6^.

Brain waste clearance has been shown to be considerably more active in sleep^7,8^. The mechanisms underlying the enhancement of waste clearance during sleep are not well understood but have been proposed to depend on an increased extracellular space during non-rapid eye movement (NREM) sleep^7^ and coupled blood-CSF flow patterns in NREM sleep^9^. Yet the importance of a complete sleep cycle, including NREM sleep, intermediate state (IS)^10^, REM sleep and awakening after each sleep cycle^11^ on vascular dynamics and CSF flow in the PVS, has not been demonstrated.

We measured the vascular dynamics throughout the sleep cycle using two-photon microscopy linescans across blood vessels in the somatosensory cortex of naturally sleeping *GLT1*-eGFP transgenic mice expressing enhanced green fluorescent protein (eGFP) in astrocytes with the vasculature outlined by intravascular Texas Red-labeled dextran (Figs. 1a,b and 2a,b)^12^. We classified sleep-wake states using an infrared sensitive camera, electrocorticography (ECoG) and electromyography (EMG) (Supplementary Fig. 1). The mice were trained to fall asleep without any use of anesthesia or sedatives, enabling us to monitor a natural progression of sleep states^12^.

**Fig. 1:**
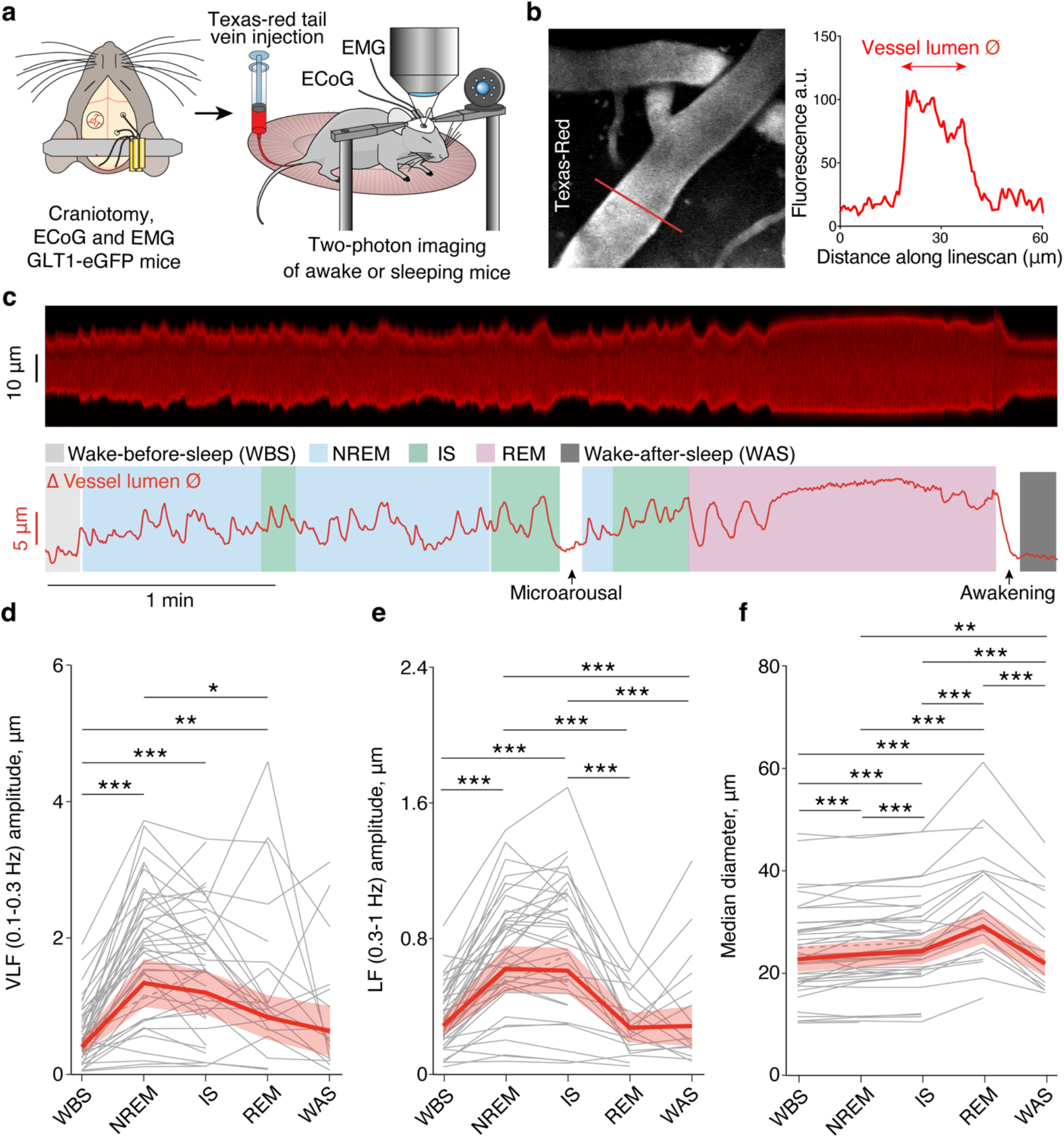
NREM, IS and REM sleep states are associated with specific pial artery diameter changes. **a,** Experimental setup. *GLT1-eGFP* transgenic mice were fitted with a cranial window exposing the somatosensory cortex, ECoG electrodes and EMG electrodes. The vasculature was outlined with Texas Red-labeled dextran. **b,** (*Left*) Representative image of a pial artery with Texas-Red labeled dextran in the lumen. Imaging was performed by placing cross sectional *x-t* line scans. (*Right*) Vessel lumen diameter was determined along a line across the vessel. **c,** Representative traces of a pial artery line scan and vessel lumen diameter during a sleep cycle. **d,** Amplitude of very low frequency (VLF 0.1–0.3 Hz), **e,** amplitude of low frequency (LF 0.3–1 Hz) oscillations and **f,** median diameter of pial artery lumen throughout a sleep cycle. Gray lines represent the individual pial arteries, dashed lines indicate missing observations in a certain state for a given artery, bold lines and shaded area are the estimates and 95% CI from linear mixed effects models. **d:** n = 570, in 44 vessels, and 5 mice, **e** and **f**: n = 579 episodes, 44 vessels, 5 mice. \**P* < 0.05, \*\**P* < 0.01, \*\*\**P* < 0.001, Tukey adjustment for multiple comparisons.

**Fig. 2:**
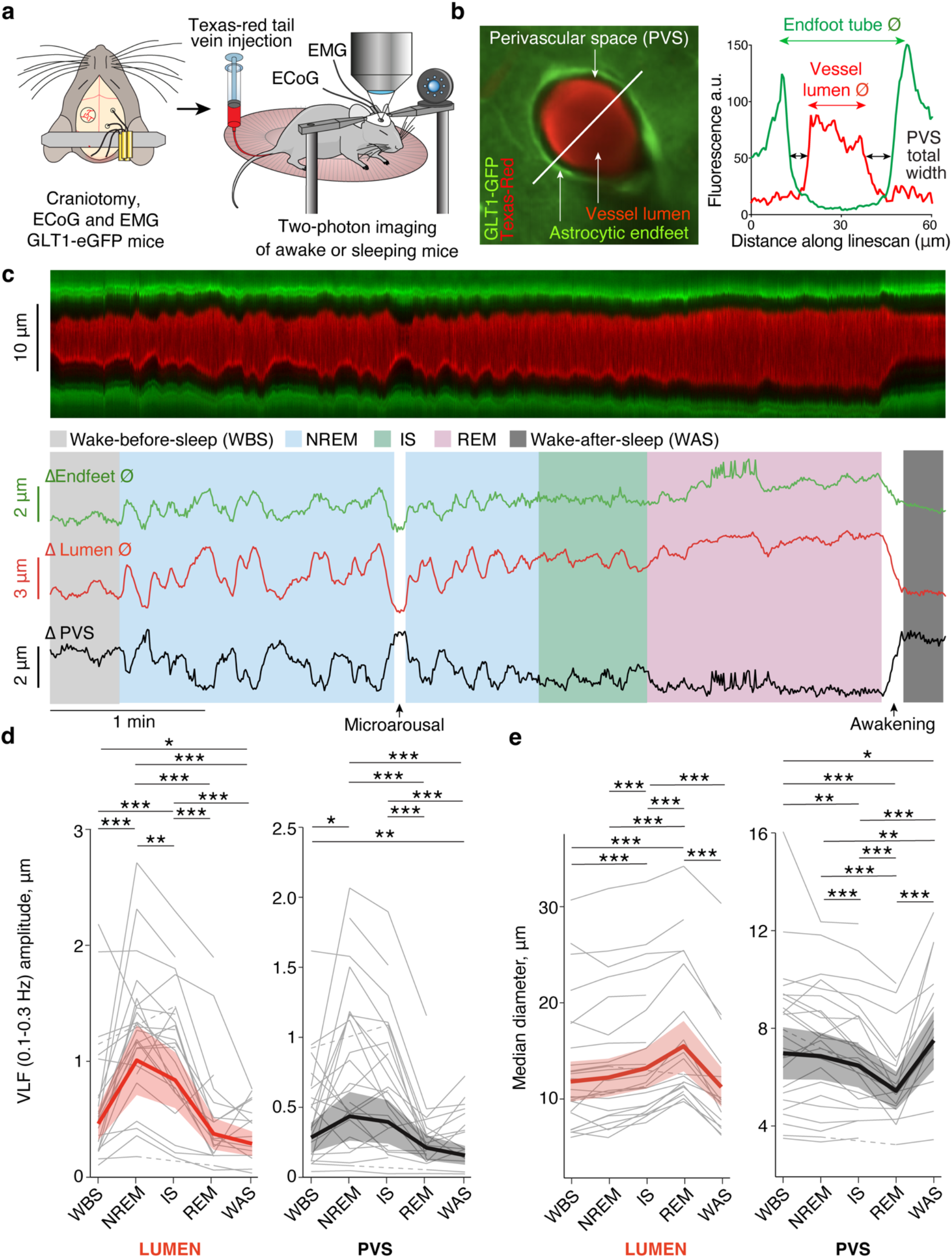
NREM, IS and REM sleep states are associated with specific penetrating arteriole diameter changes that are mirrored in the size of the PVS. **a,** Experimental setup. *GLT1*-eGFP transgenic mice were fitted with a cranial window exposing the somatosensory cortex, ECoG electrodes and EMG electrodes. The vasculature was outlined with Texas Red-labeled dextran. **b,** (*Left*) Representative image of a penetrating arteriole with *GLT1-eGFP* fluorescence in astrocytic endfeet and Texas-Red labeled dextran in the vessel lumen. Imaging was performed by placing cross sectional *x-t* line scans. (*Right*) Vessel lumen diameter and astrocyte endfoot tube diameter was determined along a line across the vessel. Total width of the PVS was assessed as the difference between the vessel lumen diameter and the endfoot tube diameter. **c,** Representative traces of a penetrating arteriole line scan, endfoot tube diameter, vessel lumen diameter and perivascular space diameter during a sleep cycle. **d,** Amplitude of very low frequency (VLF 0.1– 0.3 Hz) oscillations and **e,** median diameter of lumen and total width of PVS throughout a sleep cycle. Gray lines represent the individual penetrating arterioles, dashed lines indicate that a particular penetrating arteriole has no observations in a certain state, bold lines and shaded area are the estimates and 95% CI from linear mixed effects models, n=310 episodes, 25 penetrating arterioles, 4 mice. \**P* < 0.05, \*\**P* < 0.01, \*\*\**P* < 0.001, Tukey adjustment for multiple comparisons.

We first addressed pial arteries and measured vessel lumen diameter using *x-t* linescans (Fig. 1b). We observed striking changes in the diameter of pial arteries across the different sleep states and wakefulness (Fig. 1c). NREM and IS sleep were associated with very low frequency (VLF 0.1–0.3Hz) and low frequency (LF 0.3–1Hz) oscillations in the vessel diameter (Fig. 1d,e and Supplementary Table 1). VLF and LF oscillations of comparable amplitudes to NREM and IS sleep were only observed during locomotion across all sleep-wake states (Supplementary Fig. 2 and Supplementary Table 2). Upon REM sleep, a pronounced dilation of the arteries ensued that even outmatched the dilation observed upon locomotion (Fig. 1f, Supplementary Fig. 2 and Supplementary Tables 1–2). Subsequently, upon awakening, pial arteries constricted to reach the diameter observed during quiet wakefulness (Fig. 1f and Supplementary Table 1). Together, our data show that every part of the sleep cycle entails unique pial artery vascular dynamics.

Next, we measured vascular dynamics of penetrating arterioles (Fig. 2). Vessel lumen and endfoot tube diameters were measured in *x-t* line scans in the red and green channels, respectively, and the total width of the PVS was assessed as the difference between the vessel lumen diameter and the endfoot tube diameter (Fig. 2b). We defined PVS as the void between the lumen and endfoot sleeve to encompass all the potential pathways of CSF and solute flow in two-photon image recordings^13^. Similar to pial arteries, penetrating arterioles displayed prominent changes in the diameter of the vessel lumen (Fig. 2c). Interestingly, we also observed changes in the diameter of the endfoot tube and PVS total width (Fig. 2c). NREM and IS sleep were associated with VLF and LF oscillations in the arteriole lumen and endfoot tube diameter that were mirrored in PVS size changes (Fig. 2d, Supplementary Figs. 3a, 4 and 5b,c, and Supplementary Tables 3–6). These slow VLF and LF oscillations were considerably less prominent in all other sleep and wake states, except for locomotion (Supplementary Fig. 6a,b,d and Supplementary Tables 7–10).

When the mice entered REM sleep, the VLF and LF oscillations diminished and arteriole lumen and endfoot tube dilated, while PVS size shrunk (Fig. 2e, Supplementary Figs. 3b, 5a and Supplementary Tables 3–6). Strikingly, the magnitude of REM associated vasodilation and PVS shrinkage was larger than what was observed during locomotion (Supplementary Fig. 6c,d and Supplementary Tables 7–10). In contrast, upon awakening at the end of a sleep cycle, arteriole lumen and endfoot tube constricted to reach a similar size as in quiet wakefulness, while the PVS enlarged (Fig. 2e, Supplementary Figs. 3b, 5a and Supplementary Tables 3–6). Interestingly, the PVS was larger during wake-after-sleep than during wake-before-sleep (Fig. 2e, Supplementary Fig. 5a and Supplementary Tables 4–5). Such NREM, IS and REM associated vascular dynamics were not detected in venules (Supplementary Fig. 8). Sleep cycle-dependent vascular dynamics were confirmed in 2D time-series imaging experiments (Supplementary Fig. 7, Supplementary Video 1). Taken together, these data show that every part of a sleep cycle is associated with specific vascular and PVS dynamics of penetrating arterioles.

Next, we modeled the effects of sleep-state specific VLF and LF oscillations on fluid flow and solute transport in PVS based on individual vessel measurements (Fig. 3). We found that the VLF and LF oscillations in NREM and IS increased CSF peak velocities to levels comparable to CSF velocities measured by imaging fluorescent microbeads moving alongside pial arteries in mice^5^ and to oscillatory fluid flow generated by cardiac oscillations^14^ (Fig. 3c and Supplementary Fig. 9a), underscoring a likely salient role of slow vasomotion as a driving force for generating bulk flow in the PVS. The effect of the resultant oscillating flow on solute transport was then modeled by considering the spread of 70 kDa and 2000 kDa dextran tracer in the PVS (Fig. 3b). Compared to diffusion, VLF and LF oscillations enhanced tracer spread by dispersion, with the largest effect observed in NREM sleep (Fig. 3d,e and Supplementary Fig. 9b,c). The enhancement of VLF and LF oscillations during NREM sleep was of similar magnitude as that of cardiac oscillations (Supplementary Figs. 10a,b and 11a,b). We next analyzed how VLF and LF oscillations affected solute movement from the subarachnoid space (SAS) into the PVS of a penetrating arteriole, a process relevant not only to the glymphatic system, but also to drug delivery into the brain. VLF and LF oscillations considerably enhanced the movement of solutes from SAS into the PVS compared to pure diffusion (Fig. 3f), and consequently the solutes moved faster in NREM sleep compared to quiet wakefulness. To conclude, our modeling data suggest that VLF and LF oscillations during NREM sleep enhance CSF flow and solute transport within the arteriole PVS to levels comparable to enhancement driven by cardiac oscillations.

**Fig. 3:**
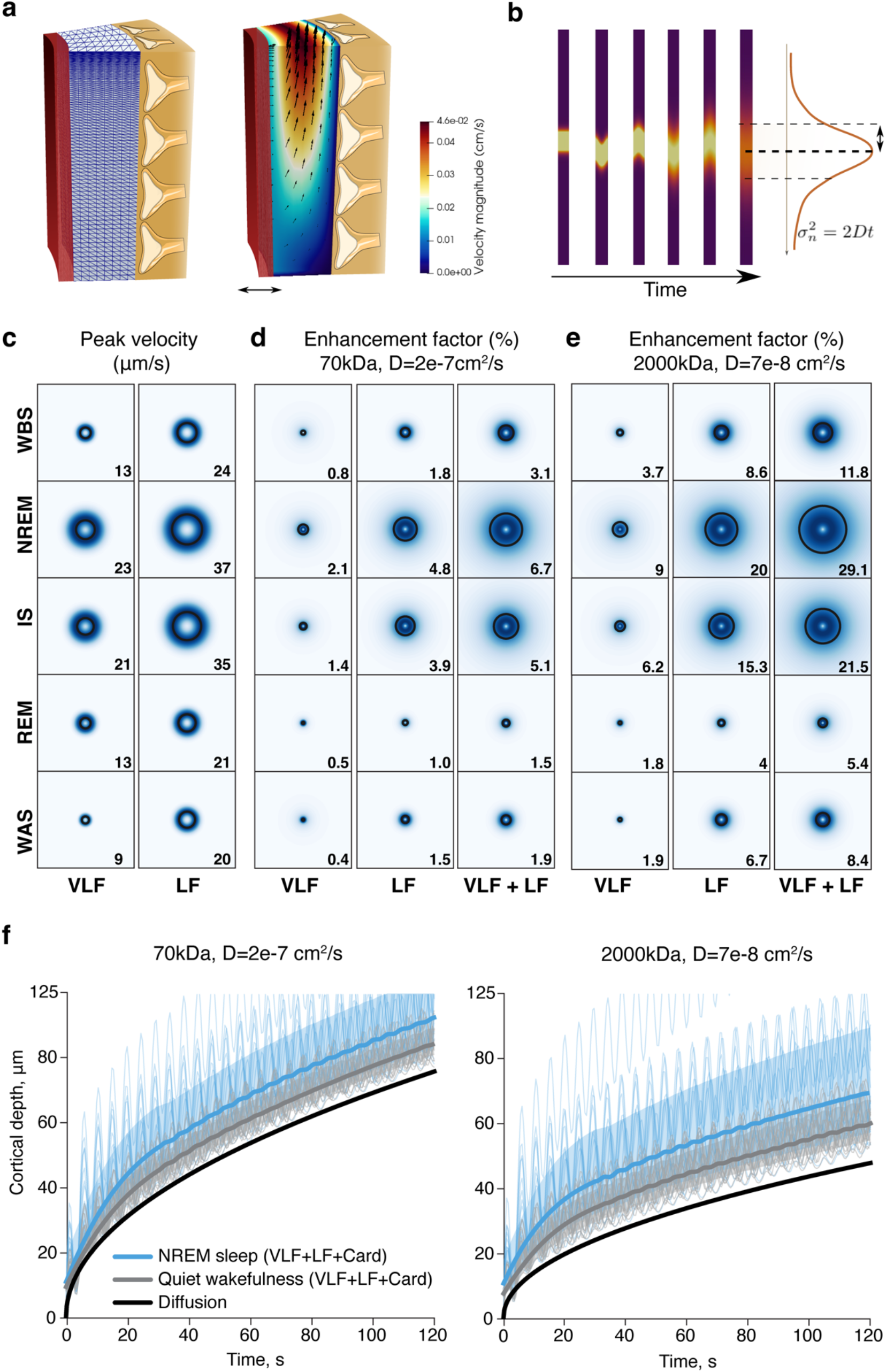
VLF and LF oscillations during NREM and IS sleep enhance CSF flow and solute transport in PVS. **a,** Illustration showing the model of the PVS (left), and representative vasomotion-driven CSF flow (right). **b,** Solute transport was assessed by the spread of tracer in the PVS. **c,** Peak velocity, **d,** enhancement factor for 70 kDa solutes, and **e,** enhancement factor for 2000 kDa solutes generated by VLF and LF oscillations during a sleep cycle as predicted by biomechanical modeling. The black circle represents the median, whereas the shading represents the distribution of all modeled vessels. The enhancement factor is the relative increase in solute movement induced by VLF and LF oscillatory flow, compared to pure diffusion. **f,** Transport of 70 kDa and 2000 kDa solutes from SAS into the PVS of penetrating arterioles driven by dispersion during quiet wakefulness, dispersion during NREM sleep or pure diffusion as predicted by biomechanical modeling. The model used the measurements of PVS cross section area for LF and VLF oscillations and assumed a peak CSF velocity of 50 μm/s for the cardiac oscillations. Thin lines represent observations from individual penetrating arterioles (n = 16 vessels, 4 mice). Bolded lines with the shading are median values of the time-smoothed thin lines over several oscillations with 10^th^ and 90^th^ percentiles. WBS - wake before sleep; WAS - wake after sleep.

The mechanisms underlying the fluid dynamics and solute transport during sleep remain elusive, partly because of the lack of data on the entire sleep cycle, including the natural progression of NREM sleep, IS sleep, REM sleep and the awakening at the end of the sleep cycle. Using naturally sleeping head-fixed mice we show that each state of the sleep cycle displays unique arteriole diameter changes that are coupled to changes in the size of PVS. By biomechanical modeling we demonstrate that slow, large-amplitude oscillatory vasomotor patterns in NREM sleep are able to generate oscillatory fluid movement on a similar magnitude to heartbeat generated fluid movement in the PVS and enhance solute transport in PVS. This supports the hypothesis that blood volume and CSF dynamics are coupled and that vasomotion could be the main driver for CSF bulk flow in the PVS^5,6,9^. The next steps will be to understand how REM sleep specific reduction in the size of PVS and the subsequent enlargement of PVS upon awakening at the end of the sleep cycle affect fluid flow and solute transport in PVS. As CSF flow has been shown to be reversed during brain-wide hyperemic patterns in NREM sleep^9^, it is likely that the REM sleep-associated vessel dilation and PVS shrinkage would drive CSF out of the brain. Conversely, during awakenings immediately after REM sleep when the vessel constricts and PVS enlarges, CSF would flow into the brain. Such a direct coupling between CSF flow direction and vascular diameter has been shown in a mouse model of ischemia, where vasoconstriction upon spreading depolarization caused a large influx of CSF into the brain^15^. Overall, we hypothesize that the entire sleep cycle is required for efficient fluid exchange and solute transport with each sleep state playing a different role in the process.

Future studies should address 3 main questions arising from this study. First, whether sleep cycle dependent vascular dynamics are global throughout the brain and whether they are synchronous along the vascular tree, or propagate in a proximal-to-distal direction or vice versa. This could depend on the type of arteriole PVS^16^. Second, how sleep cycle-dependent vascular dynamics are regulated. For instance, one potential effector for vascular dynamics could be fluctuations in norepinephrine levels across the sleep cycle^17^ and its interplay with astrocytic endfeet, that were recently shown to generate ultra-slow arteriole oscillations in awake mice^18^. Third, how vascular dynamics affect intraparenchymal waste clearance.

## METHODS

### Animals

*Glt1*-eGFP mice^19^ were housed on a 12-h light/dark cycle (lights on at 8 AM), 1–4 mice per cage. Each animal underwent surgery at the age of 8–10 weeks, followed by accommodation to be head-restrained, and two-photon imaging 2–3 times per week for up to 2 months. Adequate measures were taken to minimize pain and discomfort. Sample sizes were determined based on our previous studies using similar techniques (no power calculations were performed). All procedures were approved by the Norwegian Food Safety Authority (project number: 11983 and 22187).

### Surgical procedures

Mice were anesthetized with isoflurane. Two silver wires (200 μm thickness, non-insulated, GoodFellow) were inserted epidurally into 2 burr holes overlying the right parietal hemisphere for ECoG recordings, and two stainless steel wires (50 μm thickness, insulated except 1 mm tip, GoodFellow) were implanted in the nuchal muscles for EMG recordings. The skull over the left hemisphere was thinned for intrinsic signal imaging, a custom-made titanium head-bar was glued to the skull and the implant sealed with a dental cement cap. After two days, representations of individual whiskers in the barrel cortex were mapped by intrinsic optical imaging. The brain region activated by single whisker deflection (10 Hz, 6 s) was identified by increased red light absorption. After two days, chronic window implantation was performed. A round craniotomy of 2.5 mm diameter was made over the barrel cortex using the intrinsic optical imaging map as a reference. A window made of 2 circular coverslips of 2.5 and 3.5 mm was glued together by ultraviolet curing glue, was then centered in the craniotomy and fastened by dental cement. Mice with implant complications were excluded from the study.

### Behavioral training

Mice were housed in an enriched environment with a freely spinning wheel in their home cages. One week before imaging, mice were habituated to be head-fixed on a freely spinning wheel under the two-photon microscope. Each mouse was trained head-fixed daily before the imaging for increasing durations ranging from 10 min on the first day to 70 min on the last. Mice that showed signs of stress and did not accommodate to being head-restrained were not included in the study.

### *In vivo* two-photon imaging

Images and linescans were recorded in ScanImage (Vidrio Technologies) using a custom-built two-photon microscope (Independent NeuroScience Services) equipped with a MaiTai DeepSee ePH DS (Spectra-Physics) laser. A Nikon 16x 0.8 NA water immersion objective (model CFI75 LWD 16XW) and an excitation wavelength of 920 nm was used for imaging. Linescans were sampled at 250 Hz (149 trials) and 333 Hz (49 trials). Excitation wavelength of 920 nm was used to capture images (512 x 512 pixels) at 30 Hz in the most superficial layer for pial arteries and layer II/III for penetrating arterioles and venules of the barrel cortex. Emitted light was routed by a 565 nm longpass dichroic mirror through a 510/80 nm bandpass filter (green channel) or a 630/75 nm bandpass filter (red channel), respectively, and detected with GaAsP amplified photomultiplier tubes from Thorlabs (PMT2101). The vasculature was outlined by 2% 70 kDa Texas Red-labeled dextran (Thermo Fisher Scientific) in saline injected through a chronic tail vein catheter (200 μl at the start or imaging and 50 μl after 5 h to ensure sufficient fluorescence throughout the entire experiment). Pial arteries, arterioles and venules were distinguished based on anatomy and blood flow direction. Head-fixed sleep protocol is described in detail in Bojarskaite et al^12^. The imaging sessions of sleeping mice started at 9–10 a.m. (ZT 1–2) and lasted until 3–6 p.m. (ZT 7–10). The mice were allowed to freely move on a disc-shaped wheel/treadmill while awake. Once falling asleep the stage was locked to provide a stable platform for sleep. The position of the disc was adjusted to assist sleep in a head-fixed position. Mice that did not show any signs of sleep within the first 2 h of head-fixation were removed from the microscope. Mice were not sleep deprived or manipulated in any other way before imaging to induce sleep.

### Electrophysiological and behavioral data acquisition

ECoG and EMG signals were recorded using a DAM50 (WPI) amplifier, denoised by HumBug Noise Eliminator (Quest Scientific) and digitized by a National Instruments data acquisition device (PCIe-6351). Mouse behavior was recorded by an infrared-sensitive surveillance camera and running wheel motion. Experiments were triggered and synchronized by custom-written LabVIEW software (National Instruments).

### Sleep-wake state scoring

Wakefulness states were identified using the IR-sensitive surveillance camera video by drawing ROIs over the mouse snout and speed of the running wheel (Supplementary Fig. 1a). The signal in the snout ROIs was quantified by calculating the mean absolute pixel difference between consecutive frames in the respective ROIs. Voluntary locomotion was identified as signals above a threshold in the wheel speed time series. Spontaneous whisking was defined in the snout ROI. Quiet wakefulness was defined as wakefulness with no signal above-threshold in snout ROIs and in the wheel speed time series. Sleep states were identified from filtered ECoG (0.5–30 Hz) and EMG signals (100–1000 Hz) based on standard criteria for rodent sleep^10,20^ (Supplementary Fig. 1b): NREM sleep was defined as high-amplitude delta (0.5–4 Hz) ECoG activity and low EMG activity; IS was defined as an increase in theta (5–9 Hz) and sigma (9–16 Hz) ECoG activity, and a concomitant decrease in delta ECoG activity; REM sleep was defined as low-amplitude theta ECoG activity with theta/delta ratio >0.5 and low EMG activity. The wakefulness-before-sleep (WBS) episodes were marked as ~15 s episodes of behavioral quiescence right before NREM sleep. The wakefulness-after-sleep (WAS) episodes were marked as 10 s episodes starting immediately after awakening from REM sleep when the vessel lumen diameter stabilized. WAS typically contains mouse movement (locomotion, twitching and grooming).

### Lumen diameter, endfoot tube diameter and PVS width extraction

All data were managed with a MATLAB-based data management and analysis toolbox Begonia^21^. Line scans were recorded across penetrating arterioles, pial arteries, and veins. The linescans were cropped to center the vessel in the recordings and trials with insufficient signal quality were excluded. To improve signal quality the *x-t* data were spatiotemporally downsampled by averaging an integer number of samples to most closely reach a sampling frequency of 100 Hz and 20 samples per micrometer. To detect the radius of the lumen and the endfoot tube we created a custom MATLAB tool to manually adjust thresholds throughout the scan with a live preview for each trial to ensure an accurate tracing of both compartments. The manually specified thresholds were linearly interpolated between the chosen threshold-time point pairs. The PVS thickness was calculated by subtracting the lumen diameter from the endfoot tube diameter (Fig. 2b). The endfoot diameter, lumen diameter and PVS distance on each side of the vessel along with the specific sleep and wake states at each frame were exported to CSV files for further analysis.

### Lumen diameter, endfoot tube diameter and PVS width frequency analyses

High frequency noise was removed using the Savitzky-Golay filter with a time window of 0.1 s and a polynomial fit order of 3. The signal was decomposed into five frequency bands using Butterworth bandpass filters of order 3: continuous (0–0.1Hz), very low frequency (VLF 0.1– 0.3Hz), low frequency (LF 0.3–1Hz), respiratory (1–4Hz) and cardiac (4–15Hz) (Supplementary Fig. 12a,b). In each frequency band, local minima and maxima were detected in order to identify each individual oscillation (Supplementary Fig. 12c). The signal difference between two consecutive local minima and maxima is referred to as the peak-to-trough (P–T) amplitude. This value then represents the amplitude change in vessel diameter. The time difference between two consecutive local maxima is referred to as the P–P period.

The amplitude of cardiac and respiratory oscillations in lumen of pial arteries and penetrating arterioles are shown in Supplementary Figs. 13 and 14. The period of cardiac lumen oscillations was used for mathematical modeling described below. The amplitude for endfoot tube and PVS of penetrating arterioles in respiratory and cardiac frequencies (data not shown) was below the resolution limit of our recordings and therefore not used for further mathematical modeling analyses.

### Fluid dynamics and solute transport analysis

From the statistical analysis of the P–T amplitude, P–P period and median radius, we performed computer simulation to predict the fluid flow and solute transport in the PVS. The PVS was modeled as the space between two concentric cylinders of length 600 μm, using polar coordinates in 2D where pulsations were induced as radial changes of the inner radius. The CSF was assumed as a Newtonian, incompressible fluid with constant viscosity. The flow was described using Stokes equations and the mass transport was described using the advection diffusion equations:

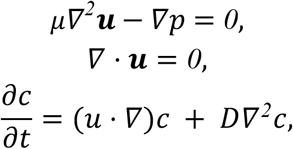

posed in the axisymmetric time-dependent domain representing the model PVS. Here, u, p and c are the fluid velocity, pressure and tracer concentration we are predicting. The fluid viscosity was taken as water viscosity at 35 °C, *μ = 0.693* mPa/s. The coefficient of diffusion, D, depends on the molecular size. For Dextran 70 kDa, the apparent diffusion *D*\* = *Dλ*^2^ measured in brain neuropil is 0.84 10^−7^ cm^2^/s ^22^. Assuming the typical value of a tortuosity *λ*=2 leads to the molecular diffusion D=1.7 × 10^−7^ cm^2^/s. For Dextran 2000 kDa we considered the molecular diffusion coefficient to be *D*=6.8× 10^−7^ cm^2^/s ^23^.

The radius of the internal cylinder *R_v_*(*t*) is assumed to be uniform along the vessel and dependent on time only. The radius of the external cylinder *R_ast_* is assumed to be fixed. We assume that both the vessel wall and the astrocyte endfoot tube are impermeable and impose a no flow condition across the walls. The variation of the cross-section area of the PVS 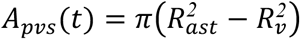 is deduced from the imaging data. It has the form 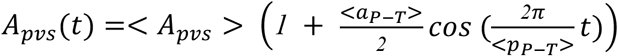 with < *A_pvs_* > and < *a*_*P–T*_ > being the median of PVS cross section area and median of the P-T amplitude of the cross section area oscillation respectively, and < *p_P–P_* > being the median of the lumen P–P period (Supplementary Figs. 14 and 15) for a given vessel, frequency band and sleep stage. We then impose the corresponding internal cylinder radius *R_v_*(*t*) and fluid velocity on the lumen wall *u_y_* = *dR_v_/dt* in the simulation. The fluid domain geometry is updated accordingly at each time step. We impose a reference pressure *p* = 0 at the entrance of the PVS (SAS side) and a no flow condition *u* = (*0,0*) at the inner side of the brain.

The equations were discretized using the finite element method in space and an implicit backward Euler scheme in time. The simulations were performed with a maximal time step of *5 · 10^−3^ s* and a maximal cell diagonal size of 1 μm.

The predicted pressure differences along the arteriole PVS are 3 Pa and 5 Pa in quiet wakefulness, and 5 Pa and 7 Pa in NREM state for VLF and LF, respectively (Supplementary Fig. 9d). The pressure gradient 7 Pa / 0.6 mm = 11.7 Pa/mm is 13.4 times larger than the estimated pressure gradients of pial PVS (0.8 Pa/mm^24^) and 11.4 times less than the upper estimate of interstitial pressure gradients (133 Pa/mm^26^).

In the dispersion analysis series (Fig. 3b,d,e), the objective was to estimate the apparent diffusion coefficient by fitting the analytical solution of the 1D pure diffusion equation. We considered the case of the diffusion of an initial Dirac delta distribution of the concentration in the middle of the PVS. The analytical solution has the form

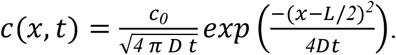

We chose 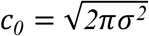 and assumed that the initial Dirac delta distribution was set at the time 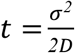. The concentration profile used at time t=0 in our simulation has therefore the form

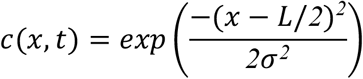

with *σ*=2 *μ*m being the standard deviation of the Gaussian profile. We then assumed that the concentration profile when the vessel wall velocity is zero remains a gaussian profile and follows the pure diffusion analytical solution. The value of the apparent diffusion *D_eff_* is determined by fitting the analytical solution to the simulation results at t =40 s.

Previous modeling studies have predicted that cardiac oscillations lead to a 100% enhancement of the transport of 70 kDa dextran by dispersion and peak oscillatory CSF flow velocity of 100–150 μm/s^14,27,28^. However, these studies deduced the PVS cross area change from the vessel wall movement only, assuming the astrocyte endfoot tube as rigid, which is not the case in our recordings (Fig. 2, Supplementary Fig. 3). The cardiac driven CSF peak velocities, when astrocyte deformation is taken into account, is expected to be much lower, on the order of 25–50 μm/s for a 250 μm long PVS^14^. The dispersion enhancement factor for such cardiac velocities is of the same order of magnitude as to what we find for the VLF and LF oscillations during NREM sleep (Fig. 3 and Supplementary Figs. 9–11). In the present study, we were only able to assess PVS size dynamics for the oscillations in the LF and VLF time scale (due to lack of imaging resolution). For cardiac oscillations, we therefore modeled three scenarios with cardiac oscillations driving CSF fluid peak velocities at 10, 50 and 100 μm/s, and imposed the associated PVS cross section area change.

In the tracer transport analysis series, the concentration was initialized at 0 in the PVS and we imposed a concentration of 1 at the SAS side. We then followed the spread of the tracer with time. We defined the concentration front as the location in the PVS where the concentration had reached 0.1.

### Statistical analyses

Each response variable, for instance the PVS amplitude in the LF frequency band, was analyzed with similar statistical methods. Because of the existence of a global rigid motion of the tissues, the displacements on both sides of a single structure (vessel or endfoot tube) should have a strong positive correlation. If not, this indicates that one edge of the structure was not well detected by our data processing tool. Unrealistic observations were therefore filtered out based on the correlation coefficient (0.8 for lumen and 0.7 for endfoot tube) between the position of both sides of the structure (vessel or astrocytes endfoot tube). Then, the data were aggregated by taking the average of observations belonging to each episode. Here an episode is defined as a continuous time period within a trial where the mouse is identified as behaving according to a single sleep-wake state. A linear mixed effect model was then fitted to the aggregated data set, after log transforming the response variable. For each response variable a sequence of successively simpler models was attempted until reaching numerical convergence and satisfactory looking residual plots. All candidate models had sleep-wake state as a categorical fixed effect, with quiet wakefulness as the baseline state (i.e., the intercept). The models for amplitude and period also included the median lumen radius as a continuous second-order fixed effect, as long as there was sufficient data to estimate these effects. For some response variables, for example for lumen LF amplitude, there was a significant relationship between the response (here amplitude) and median lumen radius. This relationship took different forms, but could often take the form of an inverted U shape, with high expected amplitude for middling median lumen radii, and lower expected amplitude for both small and high lumen radii. All candidate models included random intercepts for each vessel (or penetrating arteriole). For most responses there was large variation between vessels (as seen for example in the grey lines in Fig 1d,e.). For response variables where more complex models could be fit (due to sufficient data and depending on residual plots) we also included random effects on the state effect, meaning that each vessel could potentially have somewhat different state-to-state differences. Further, the more complex models also include the possibility of heterogeneous residual variance, in each sleep-wake state and for each mouse. After model fitting, pairwise contrasts between each state estimate were computed. For each fitted model, the p-values belonging to the multiple resulting contrasts were adjusted by Tukey’s method for multiple comparisons. Statistical analyses were conducted in R (version 4.0.5). The linear mixed effect models were fitted using the glmmTMB package^29^, residual plots were constructed by the DHARMa package (https://CRAN.R-project.org/package=DHARMa), and contrasts computed by the emmeans package (https://CRAN.R-project.org/package=emmeans).

## Data availability

The datasets generated in this study are available upon reasonable request. Source data for the figures are provided with the manuscript.

## Code availability

Code for data management and vascular diameter extraction: https://github.com/GliaLab/PVS-Sleep-Project. Code for vascular diameter change analysis, fluid flow simulations, tracer transport simulations and dispersion analysis: https://github.com/AlexandraVallet/PVSflow. Code for statistical analyses can be found here: https://drive.google.com/drive/folders/1KlxkVbjiRIadiggpqWZmdp8zx-RG2CEh.

## Acknowledgements

This work was supported by the Medical Faculty, University of Oslo, the Olav Thon Foundation, the Letten Foundation, the Research Council of Norway (grants #249988, #271555/F20, #302326 #300305, #301013, #303362), the South-Eastern Norway Regional Health Authority (grant #2016070). We acknowledge the support by UNINETT Sigma2 AS for computational science, grant NN9279K and for making data storage available through NIRD, project NS9021K. Associate Professor Gudmund Horn Hermansen is gratefully acknowledged for his input pertaining statistical modeling.

## Author Contributions

Conceptualization: L.B., A.V., K.A.M., R.E. Methodology: L.B., A.V., D.M.B., C.C., M.K., K.A.M., R.E. Formal analysis: D.M.B., A.V., C.C. Investigation: L.B. and K.M.G.B, Visualization: L.B., A.V., C.C. Data Curation: D.M.B., A.V., C.C. Software: D.M.B., A.V., C.C., M.K. Funding acquisition: K.A.M. and R.E. Project administration: L.B., D.M.B., A.V., K.A.M., R.E. Resources: K.A.M., K.H. and R.E. Supervision: K.A.M and R.E. Writing - original draft and revision: L.B., R.E., A.V., C.C., K.A.M. Writing - review and editing: K.M.G.B., K.H., M.K., C.C., D.M.B., L.B., R.E., A.V., K.A.M.

