## Supplementary Information for "Sleep cycle-dependent vascular dynamics enhance perivascular cerebrospinal fluid flow and solute transport"

#### Supplementary data

This supplementary data contains:

Supplementary figures 1-15
Supplementary tables 1-10
Supplementary video 1

### Supplementary figures

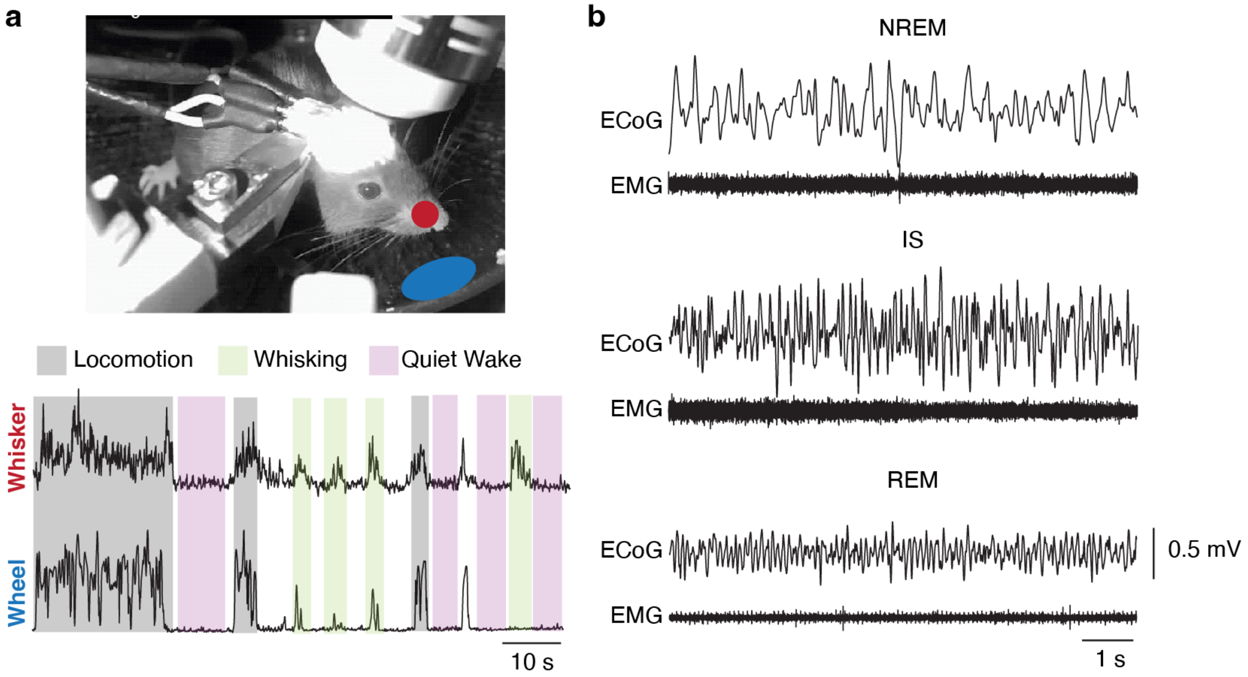


**Supplementary Fig. 1: Sleep-wake state scoring. a**, Wakefulness was separated into locomotion, whisking and quiet wakefulness based on movement detected in snout and running wheel regions in the surveillance video. **b,** Representative ECoG and EMG traces during NREM, IS and REM sleep.

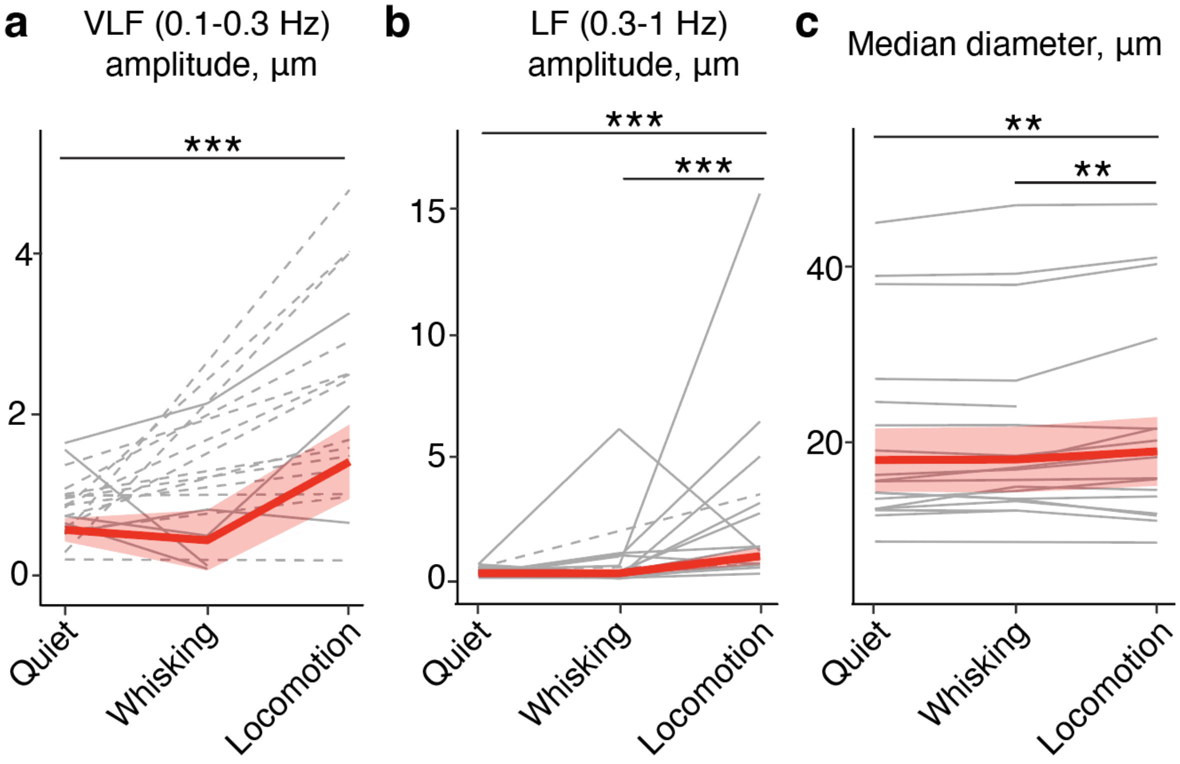


**Supplementary Fig. 2: a,** Amplitude of very low frequency (VLF 0.1–0.3 Hz) oscillations, **b,** amplitude of low frequency (LF 0.3-1 Hz) oscillations, and **c,** median diameter of lumen of pial arteries during quiet wakefulness, whisking and locomotion. Gray lines represent the individual blood vessels, dashed lines indicate that a particular penetrating arteriole has no observations in a certain state, bold lines and shaded area are the estimates and 95% CI from linear mixed effects models. For **a**: n = 498 episodes, 19 vessels, 5 mice; for **b**: n = 797 episodes, 19 vessels, 5 mice; for **c** n = 888 episodes, 19 vessels, 5 mice. *$P < 0.05$, **$P < 0.01$, ***$P < 0.001$, Tukey adjustment for multiple comparisons.

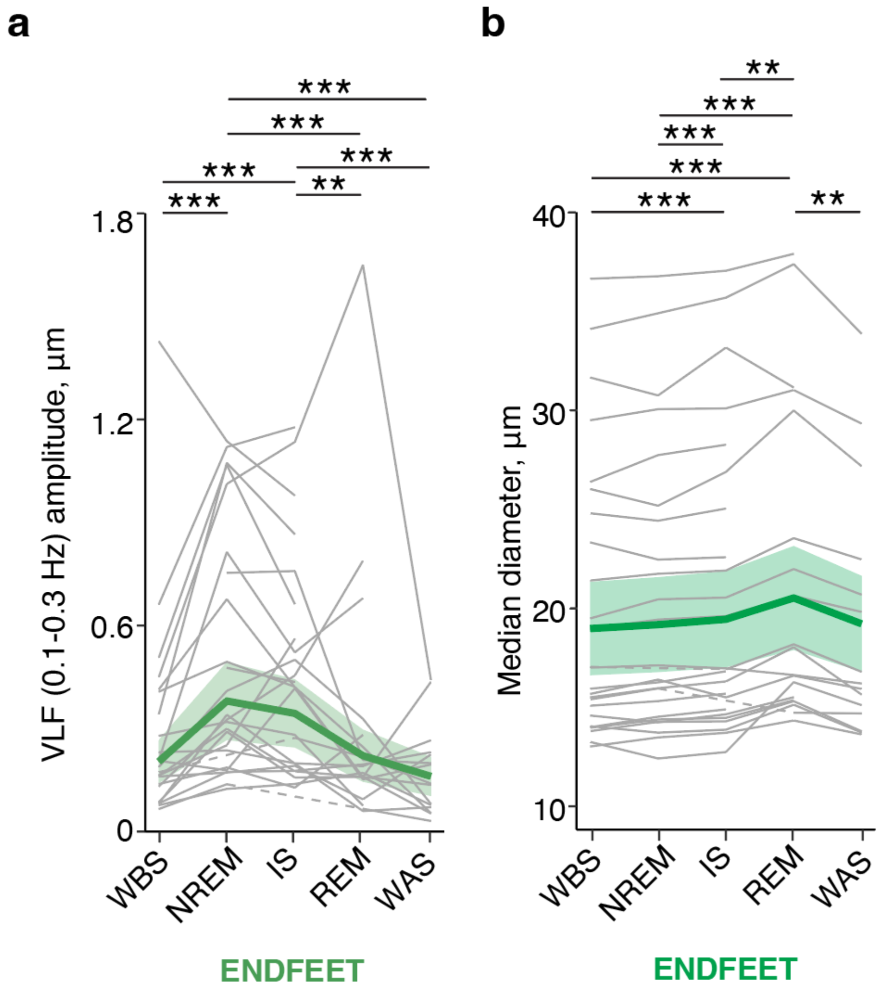


**Supplementary Fig. 3**: **a,** Amplitude of very low frequency (VLF 0.1–0.3 Hz) oscillations and **b,** median diameter of enfoot tube during a sleep cycle. Gray lines represent the individual blood vessels (median value of all episodes of the same sleep-wake state), dashed lines indicate that a particular penetrating arteriole has no observations in a certain state, bold lines and shaded area are the estimates and 95% CI from the linear mixed effects statistical model. For **a**: n = 306 episodes, 25 vessels, 4 mice; for **b**: n = 310 episodes, 25 vessels, 4 mice. *$P < 0.05$, **$P < 0.01$, ***$P < 0.001$, Tukey adjustment for multiple comparisons. WBR - wake before sleep, WAS - wake after sleep.

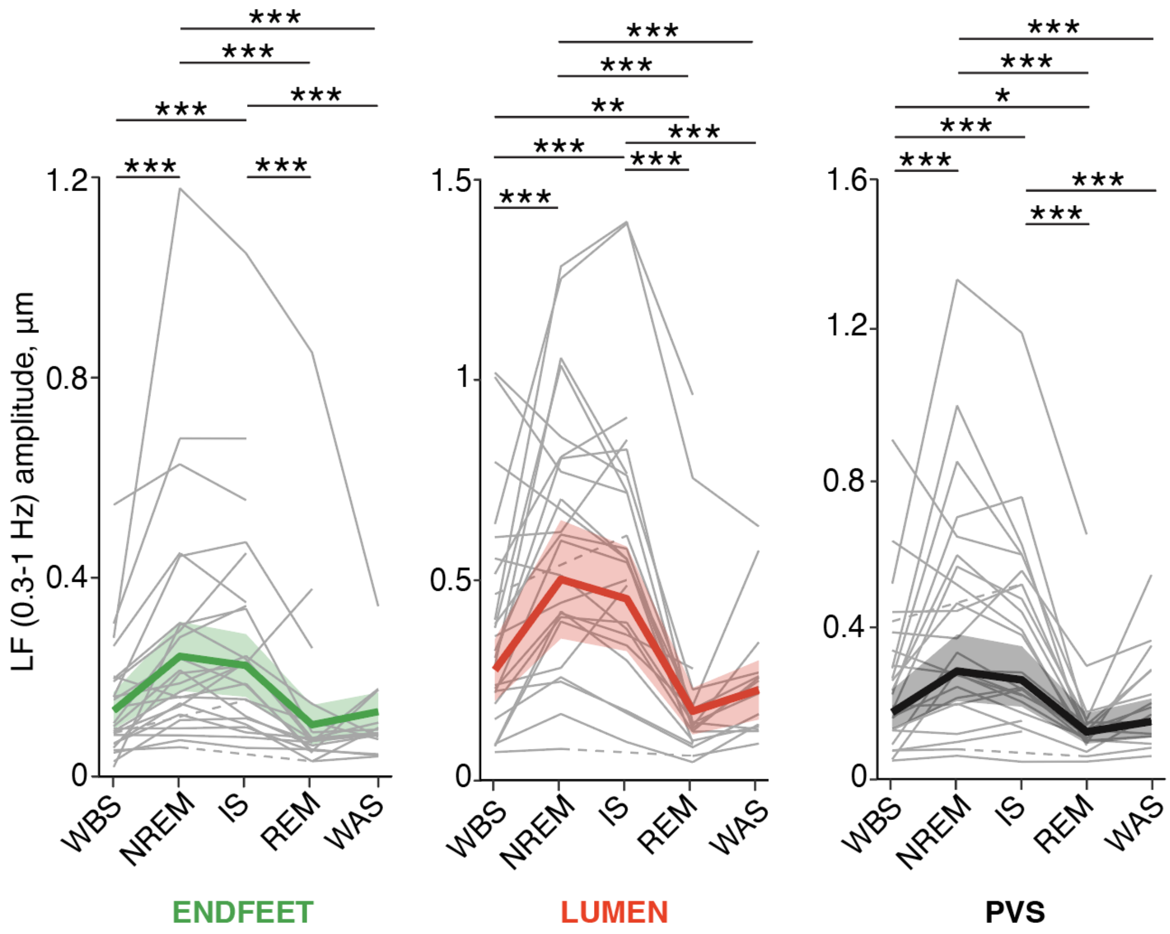


**Supplementary Fig. 4**: Amplitude of low frequency (LF 0.3–1 Hz) oscillations in endfoot tube, lumen and PVS during a sleep cycle. Gray lines represent the individual penetrating arterioles, dashed lines indicate that a particular penetrating arteriole has no observations in a certain state, bold lines and shaded area are the estimates and 95% CI from linear mixed effects models, n = 310 episodes, 25 vessels, 4 mice. *$P < 0.05$, **$P < 0.01$, ***$P < 0.001$, Tukey adjustment for multiple comparisons. WBR - wake before sleep, WAS - wake after sleep.

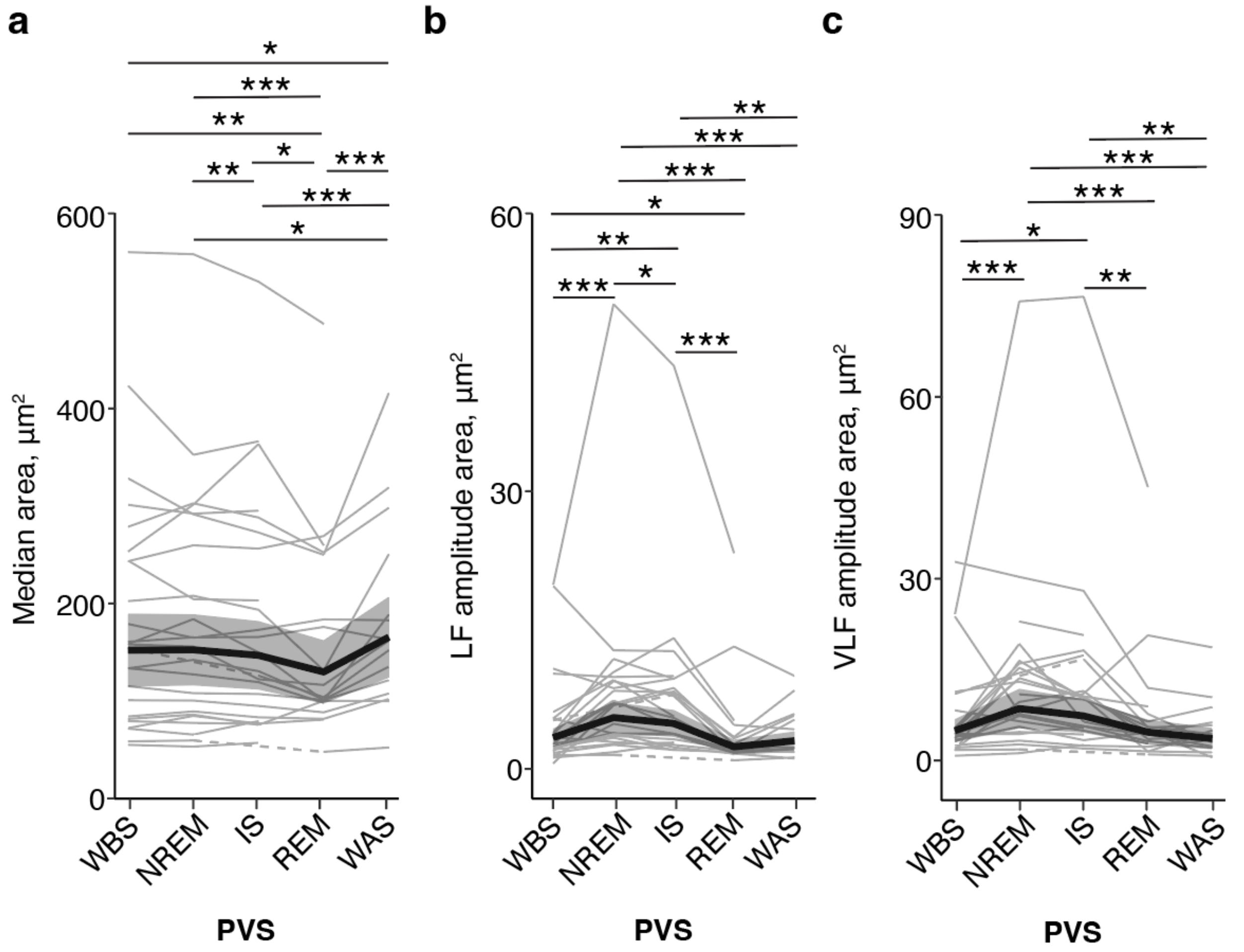


**Supplementary Fig. 5**: **a,** Median PVS area, **b,** LF amplitude in PVS area and **c,** VLF amplitude in PVS area during a sleep cycle. Gray lines represent the individual penetrating arterioles, dashed lines indicate that a particular penetrating arteriole has no observations in a certain state, bold lines and shaded area are the estimates and 95% CI from linear mixed effects models. For **a** and **b**: n = 310 episodes, 25 vessels, 4 mice; for **c**: n = 304 episodes, 25 vessels, 4 mice. *$P < 0.05$, **$P < 0.01$, ***$P < 0.001$, Tukey adjustment for multiple comparisons. WBR - wake before sleep, WAS - wake after sleep.

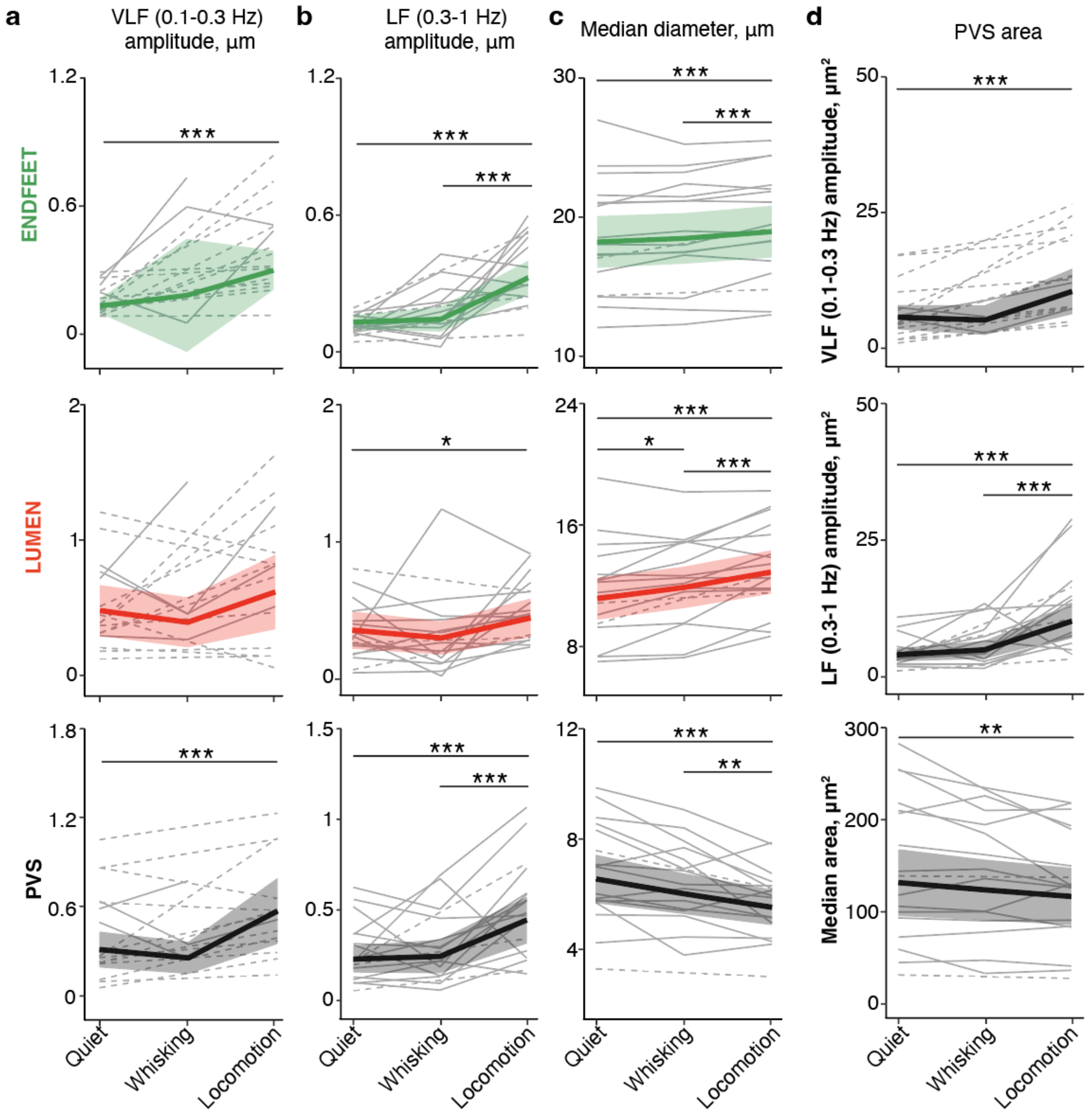


**Supplementary Fig. 6: a,** Amplitude of very low frequency (VLF 0.1–0.3 Hz) oscillations, **b,** amplitude of low frequency (LF 0.3–1 Hz) oscillations, and **c,** median diameter (width for PVS) of (*top-to-bottom*) endfoot tube, lumen and PVS of penetrating arterioles during quiet wakefulness, whisking and locomotion. **d,** (*top-to-bottom*) VLF amplitude in PVS area, LF amplitude in PVS area and median PVS area during quiet wakefulness, whisking and locomotion. Gray lines represent the individual penetrating arterioles, dashed lines indicate that a particular penetrating arteriole has no observations in a certain state, bold lines and shaded area are the estimates and 95% CI from linear mixed effects models. **a**: n = 127 episodes, 16 vessels, 3 mice for endfoot tube and lumen, n = 134 episodes, 16 vessels, 3 mice for PVS; **b:** n = 236 episodes, 17 vessels, 3 mice for endfoot tube, n = 228 episodes, 17 vessels, 3 mice for lumen, n = 244 episodes, 17 vessels, 3 mice for PVS; **c**: n = 262 episodes, 17 vessels, 3 mice **d**: n = 134 episodes, 16 vessels, 3 mice for VLF amplitude, n = 243 episodes, 17 vessels, 3 mice for LF amplitude, n = 262 episodes, 17 vessels, 3 mice for median area. *$P < 0.05$, **$P < 0.01$, ***$P < 0.001$, Tukey adjustment for multiple comparisons.

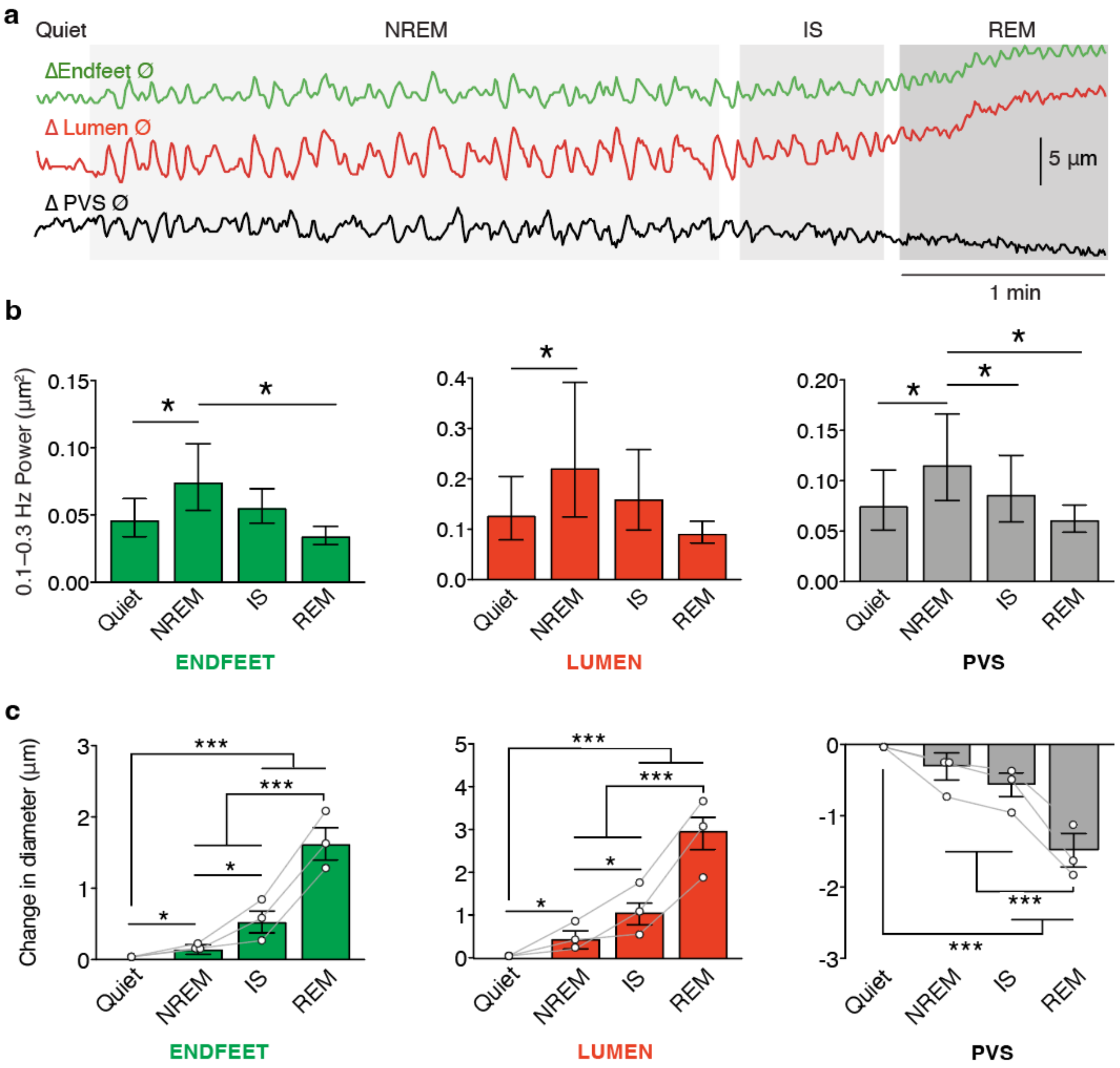


**Supplementary Fig. 7**: **2D time-series analysis of vascular dynamics across the sleep cycle. a,** Representative traces of a penetrating arteriole endfoot tube diameter, vessel lumen diameter and PVS total width during quiet wakefulness, NREM, IS and REM sleep states. **b,** Average diameter oscillation power in the 0.1–0.3 Hz frequency range of endfoot tube, vessel lumen and PVS during quiet wakefulness, NREM, IS and REM sleep. Data represented as estimates ± standard error, n = 3 mice, 72 arterioles. **c,** Change in the median diameter per sleep state of the endfoot tube, vessel lumen, and perivascular space of penetrating arterioles from quiet wakefulness to NREM, IS and REM sleep. Data represented as estimates ± SE, n = 3 mice, 136 arterioles. *$P < 0.05$, **$P < 0.01$, ***$P < 0.001$.

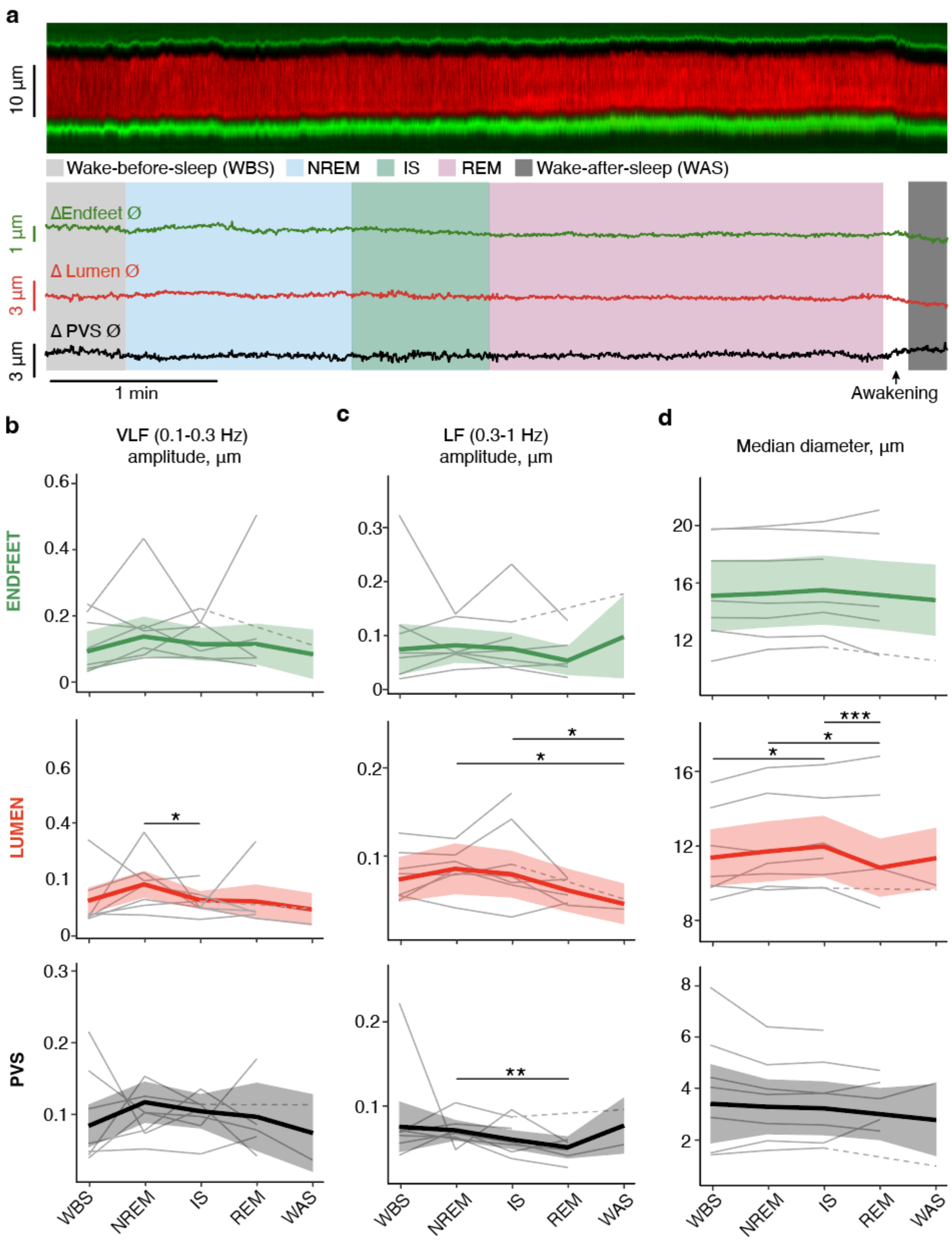


**Supplementary Fig. 8: a,** Representative traces of a venule line scan, endfoot tube diameter, vessel lumen diameter and PVS width during a sleep cycle. Gray lines represent the individual penetrating arterioles, dashed lines indicate that a particular penetrating arteriole has no observations in a certain state, bold lines and shaded area are the estimates and 95% CI from linear mixed effects statistical model, n = 90 episodes, 7 vessels, 3 mice. *$P < 0.05$, **$P < 0.01$, ***$P < 0.001$, Tukey adjustment for multiple comparisons. WBR - wake before sleep, WAS - wake after sleep.

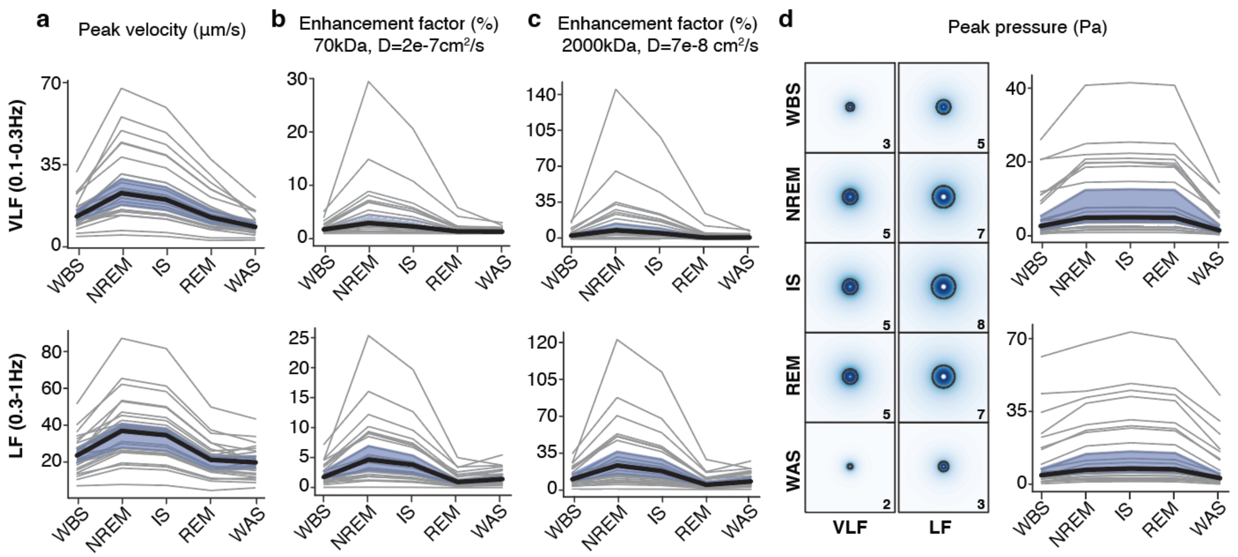


**Supplementary Fig. 9: VLF and LF oscillations during NREM and IS sleep enhance CSF flow and solute transport in PVS. a,** Peak velocity, **b,** enhancement factor for 70 kDa solutes, **c,** enhancement factor for 2000 kDa solutes and **d,** peak pressure generated by VLF and LF oscillations across states of a sleep cycle as predicted by biomechanical modeling. The blue color surface represents the distribution obtained from all observations, the black line is the median of the observations. Gray lines represent observations from individual penetrating arterioles while bolded black lines with the blue shading are median values with 10$^{th}$ and 90$^{th}$ percentiles, n = 25 vessels, 4 mice. WBS - wake before sleep; WAS - wake after sleep.

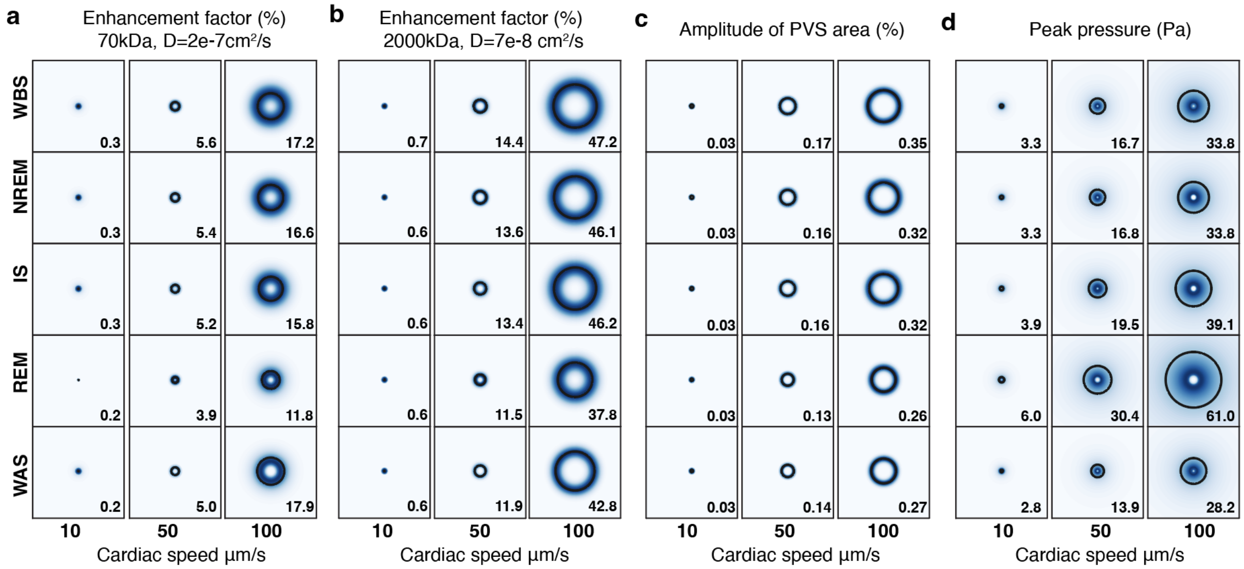


**Supplementary Fig. 10**: **Biomechanical modeling of CSF flow and solute transport in PVS generated by cardiac pulsations with different peak CSF velocities. a,** Enhancement factor for 70 kDa solutes, **b,** enhancement factor for 2000 kDa solutes, **c,** amplitude of PVS area, and **d,** peak pressure generated by cardiac pulsations with peak CSF velocity of 10, 50 or 100 µm/s across states of a sleep cycle as predicted by biomechanical modeling. The blue color surface represents the distribution obtained from all observations, the black line is the median of the observations. WBS - wake before sleep; WAS - wake after sleep.

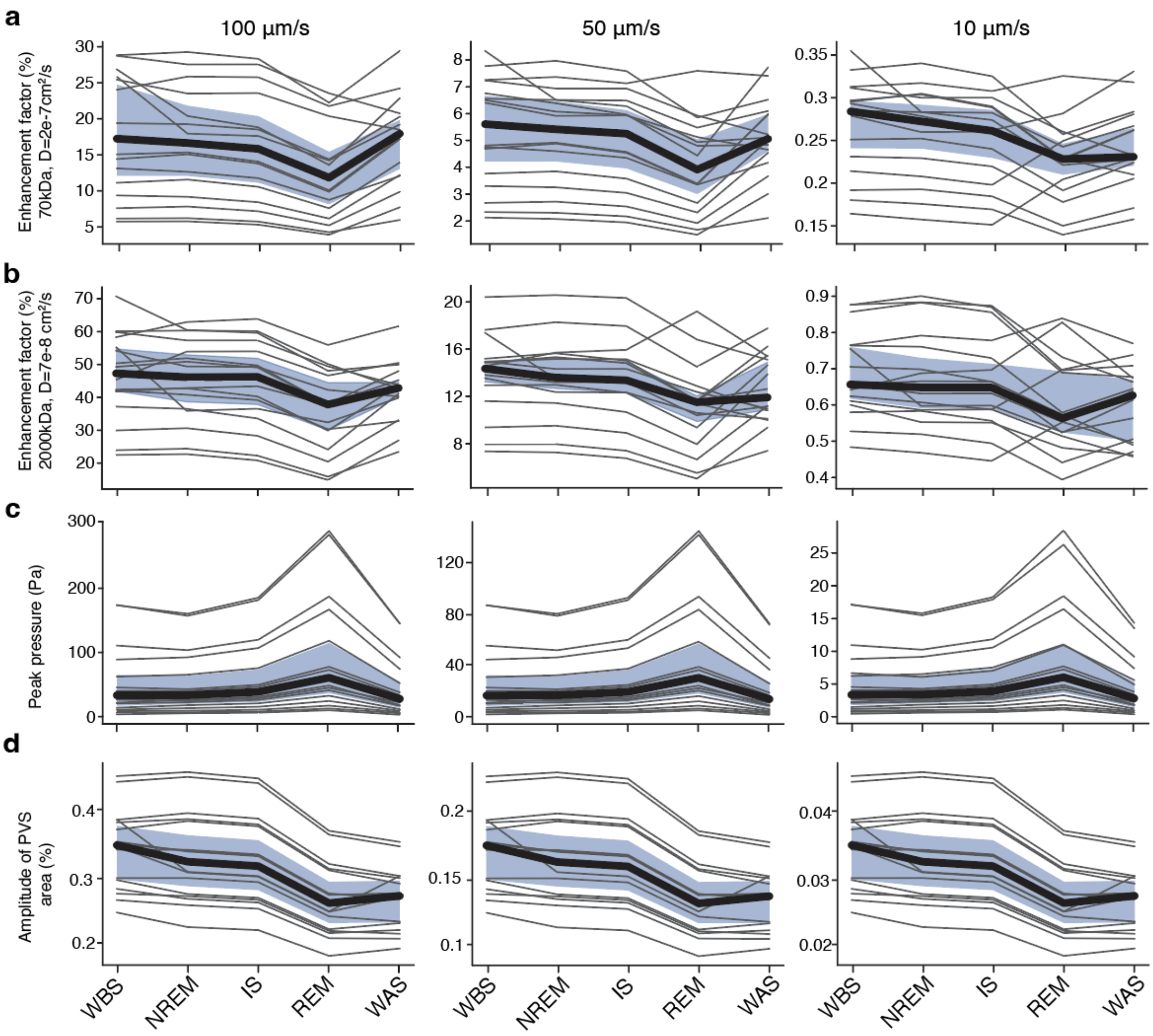


**Supplementary Fig. 11: Biomechanical modeling of CSF flow and solute transport in PVS generated by cardiac pulsations with different peak CSF velocities. a,** Enhancement factor for 70 kDa solutes, **b,** enhancement factor for 2000 kDa solutes, **c,** peak pressure**,** and **d,** amplitude of PVS area, generated by cardiac pulsations with peak CSF velocity of 10, 50 or 100 µm/s across the states of a sleep cycle as predicted by biomechanical modeling. Gray lines represent observations from individual penetrating arterioles while bolded black lines with the blue shading are median values with 10$^{th}$ and 90$^{th}$ percentiles, n = 16 vessels, 4 mice. WBS - wake before sleep; WAS - wake after sleep.

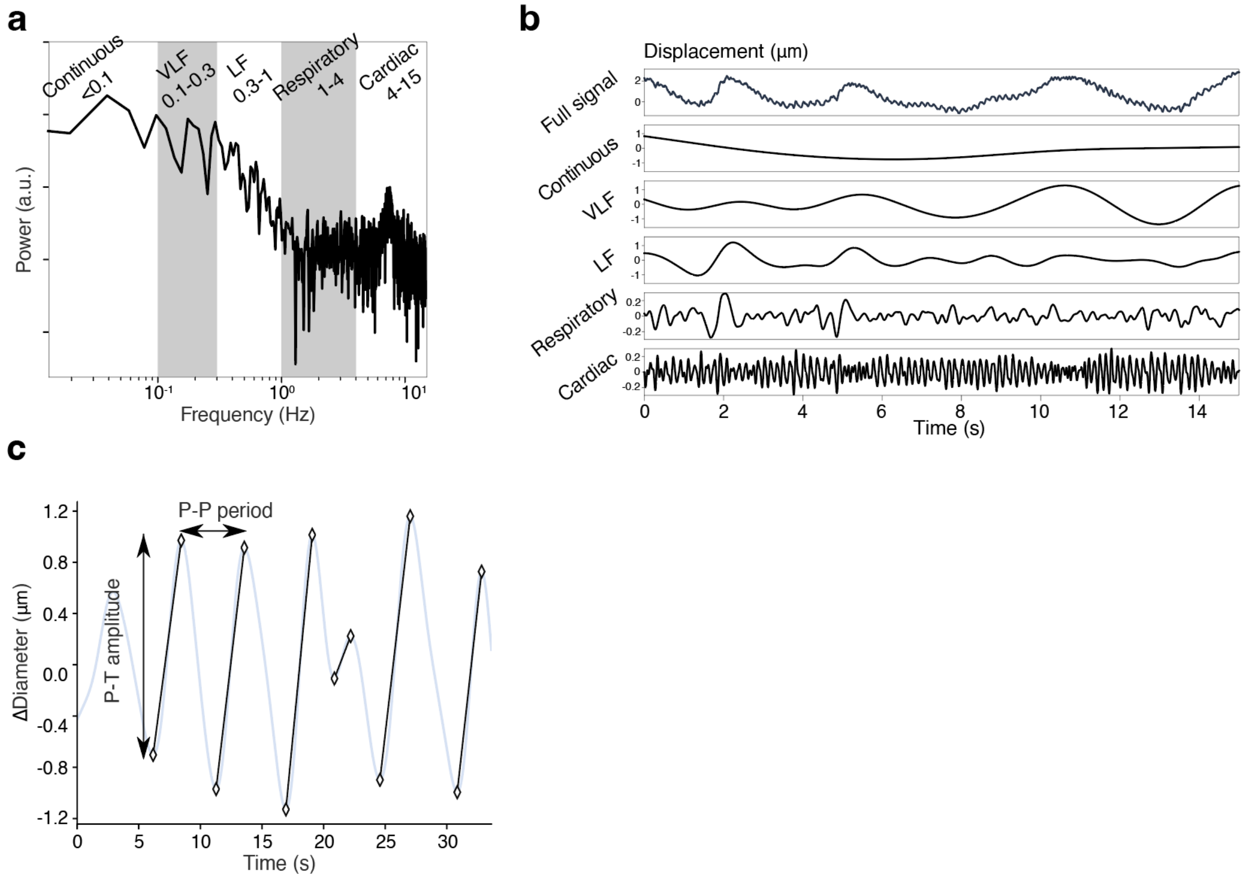


**Supplementary Fig. 12: a,** Power spectrum of vessel diameter oscillations and different frequency bands used for analysis. **b,** Signal decomposition into continuous, VLF, LF, respiratory and cardiac frequency bands. **c,** Amplitude of oscillations in different frequency bands was calculated as peak-to-trough (P-T), while period as peak-to-peak (P-P). VLF - very low frequency; LF - low frequency.

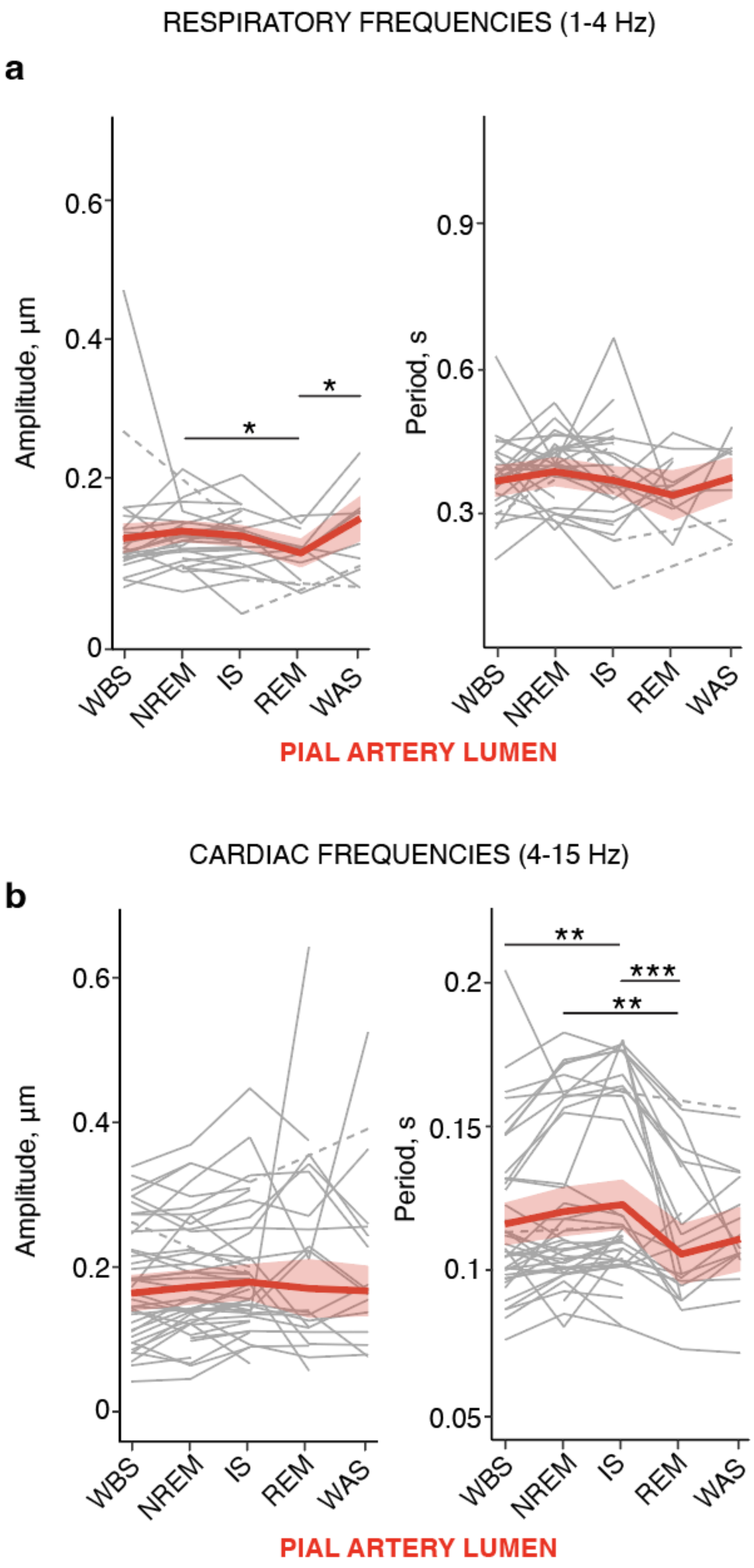
RESPIRATORY FREQUENCIES (1-4 Hz)
a
Amplitude, µm
0.6
0.4
0.2
0
*
*
Period, s
0.9
0.6
0.3
WBS
NREM
IS
REM
WAS
PIAL ARTERY LUMEN
CARDIAC FREQUENCIES (4-15 Hz)
b
Amplitude, µm
0.6
0.4
0.2
0
Period, s
0.2
0.15
0.1
0.05
**
***
**
WBS
NREM
IS
REM
WAS
PIAL ARTERY LUMEN

**Supplementary Fig. 13:** Amplitude and period of **a,** respiratory (1-4Hz) and **b,** cardiac (4-15Hz) frequencies of pial artery lumen throughout a sleep cycle. Gray lines represent the individual penetrating arterioles, dashed lines indicate that a particular penetrating arteriole has no observations in a certain state, bold lines and shaded area are the estimates and 95% CI from linear mixed effects models, n = 343 episodes, 30 pial arteries, 5 mice for **a**, n=487 episodes, 43 pial arteries, 4 mice for **b**. $*P < 0.05$, $**P < 0.01$, $***P < 0.001$, Tukey adjustment for multiple comparisons.

RESPIRATORY FREQUENCIES (1-4 Hz)

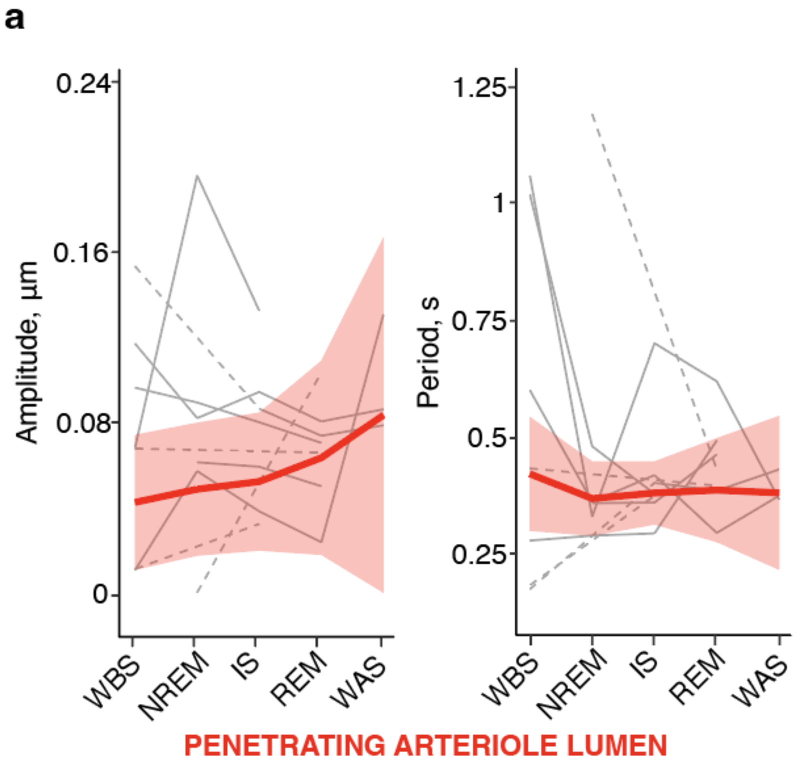


CARDIAC FREQUENCIES (4-15 Hz)

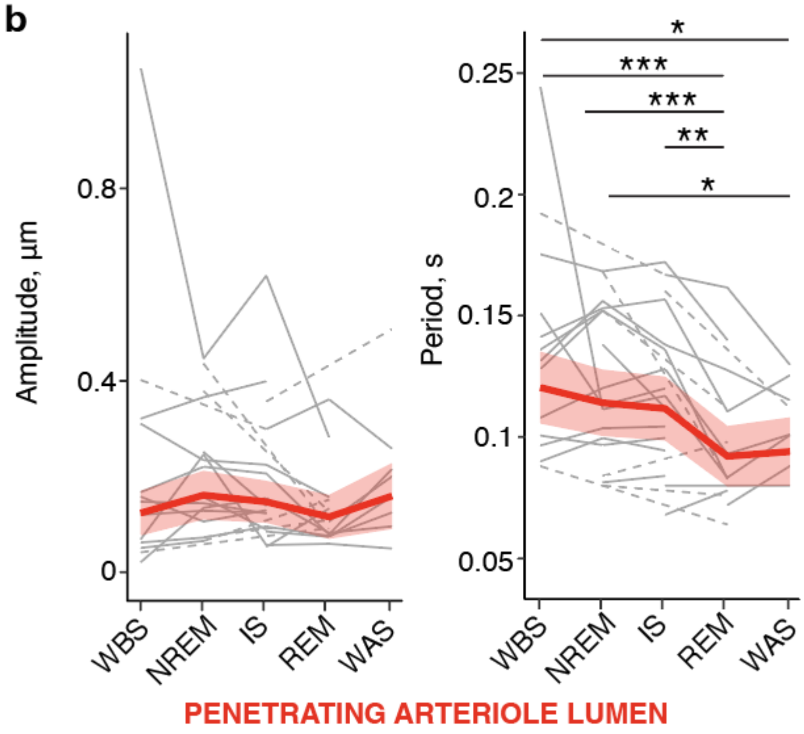

**Supplementary Fig. 14:** Amplitude and period of **a,** respiratory (1-4Hz) and **b,** cardiac (4-15Hz) frequencies of penetrating arteriole lumen throughout a sleep cycle. Gray lines represent the individual penetrating arterioles, dashed lines indicate that a particular penetrating arteriole has no observations in a certain state, bold lines and shaded area are the estimates and 95% CI from linear mixed effects models, n=135 episodes, 22 penetrating arterioles, 4 mice for **a**, n=57 episodes, 9 penetrating arterioles, 3 mice for **b**. $^{*}P < 0.05$, $^{**}P < 0.01$, $^{***}P < 0.001$, Tukey adjustment for multiple comparisons.

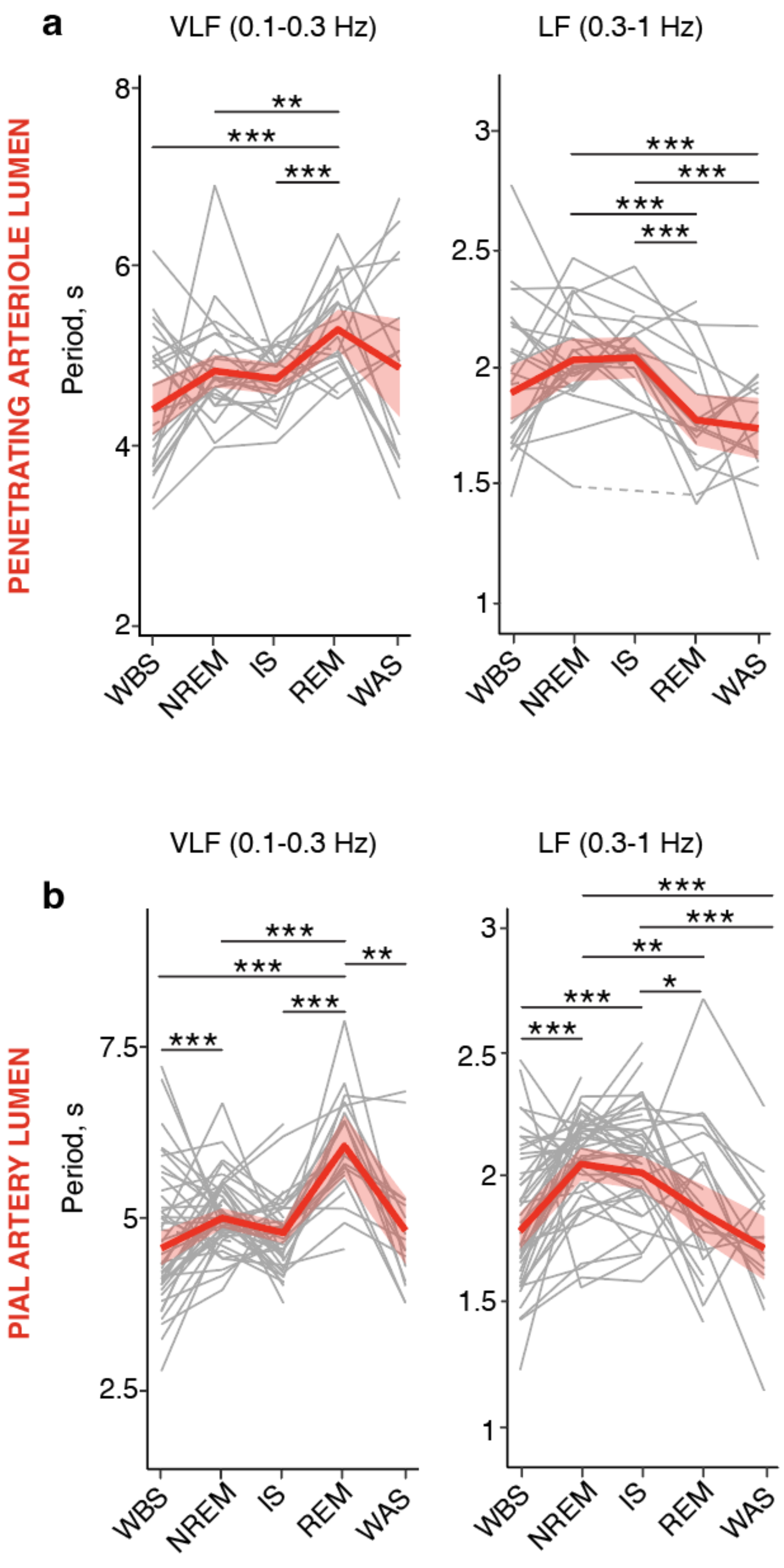

a
VLF (0.1-0.3 Hz)
LF (0.3-1 Hz)
PENETRATING ARTERIOLE LUMEN
Period, s
WBS
NREM
IS
REM
WAS
b
PIAL ARTERY LUMEN

**Supplementary Fig. 15:** Period of VLF and LF oscillations in lumen of **a,** pial arteries and **b,** penetrating arterioles throughout a sleep cycle. Gray lines represent the individual penetrating arterioles, dashed lines indicate that a particular penetrating arteriole has no observations in a certain state, bold lines and shaded area are the estimates and 95% CI from linear mixed effects models, n=310 episodes, 25 penetrating arterioles, 4 mice for **a**, n=570 episodes, 44 penetrating arterioles, 5 mice for **b**. $^{*}P < 0.05$, $^{**}P < 0.01$, $^{***}P < 0.001$, Tukey adjustment for multiple comparisons.

Supplementary tables

| **Lumen diameter** | Wake before sleep | NREM | IS | REM | Wake after sleep |
|---|---|---|---|---|---|
| VLF amplitude, µm (%) | 0.4 (1.8) | 1.4 (5.9) | 1.2 (4.9) | 0.9 (3.1) | 0.7 (3.2) |
| LF amplitude, µm (%) | 0.3 (1.3) | 0.6 (2.5) | 0.6 (2.5) | 0.3 (1.0) | 0.3 (1.4) |
| Median diameter, µm (%) | 22.7 | 23.6 (4.0) | 24.4 (7.5) | 29.2 (28.6) | 21.6(-4.8) |

**Supplementary Table 1**. Estimated values of VLF amplitude, LF amplitude and diameter of pial artery lumen during sleep states obtained from linear mixed effects models. For VLF amplitude and LF amplitude, numbers in brackets show the percentage wise change in diameter from median diameter in each sleep cycle state. For median diameter, numbers in brackets show percentage wise change in diameter from wake-before-sleep state.

| **Lumen diameter** | Quiet wakefulness | Whisking | Locomotion |
|---|---|---|---|
| VLF amplitude, µm (%) | 0.6 (3.3) | 0.4 (2.2) | 1.4 (7.4) |
| LF amplitude, µm (%) | 0.3 (1.7) | 0.3 (1.7) | 0.7 (3.7) |
| Median diameter, µm (%) | 18 | 18.1 (0.6) | 19.0 (5.6) |

**Supplementary Table 2.** Estimated values of VLF amplitude, LF amplitude and diameter of pial artery lumen during wake states obtained from linear mixed effects models. For VLF amplitude and LF amplitude, numbers in brackets show the percentage wise change in diameter from median diameter in each wakefulness state. For median diameter, numbers in brackets show percentage wise change in diameter from quiet wakefulness state.

| **Lumen diameter** | Wake before sleep | NREM | IS | REM | Wake after sleep |
|---|---|---|---|---|---|
| VLF amplitude, µm | 0.4 (3.4) | 1 (8.2) | 0.8 (6.1) | 0.4 (2.6) | 0.3 (2.7) |
| LF amplitude, µm | 0.3 (2.5) | 0.5 (4.1) | 0.45 (3.4) | 0.2 (1.3) | 0.2 (1.8) |
| Median diameter, µm | 11.9 | 12.2 (2.5) | 13.2 (10.9) | 15.5 (30.3) | 11.3 (-5.0) |

**Supplementary Table 3.** Estimated values of VLF amplitude, LF amplitude and diameter of penetrating arteriole lumen during sleep states obtained from linear mixed effects models. For VLF amplitude and LF amplitude, numbers in brackets show the percentage wise change in diameter from median diameter in each sleep cycle state. For median diameter, numbers in brackets show percentage wise change in diameter from wake-before-sleep state.

| **PVS total width** | Wake before sleep | NREM | IS | REM | Wake after sleep |
|---|---|---|---|---|---|
| VLF amplitude, µm (%) | 0.29 (4.7) | 0.44 (7.2) | 0.4 (7.1) | 0.21 (4.7) | 0.16 (2.4) |
| LF amplitude, µm (%) | 0.16 (2.6) | 0.27 (4.4) | 0.25 (4.5) | 0.11 (2.4) | 0.14 (2.1) |
| Median total width, µm (%) | 6.2 | 6.1 (-1.6) | 5.6 (-9.7) | 4.5 (-27.4) | 6.8 (9.7) |

**Supplementary Table 4**. Estimated values of VLF amplitude, LF amplitude and total width of penetrating arteriole PVS during sleep states obtained from linear mixed effects models. For VLF amplitude and LF amplitude, numbers in brackets show the percentage wise change in diameter from median diameter in each sleep cycle state. For median total width, numbers in brackets show percentage wise change in total width from wake-before-sleep state.

| **PVS area** | Wake before sleep | NREM | IS | REM | Wake after sleep |
|---|---|---|---|---|---|
| VLF amplitude, $\mu m^2$ (%) | 5 (3.4) | 8.7 (5.9) | 7.5 (5.2) | 4.8 (3.8) | 3.7 (2.3) |
| LF amplitude, $\mu m^2$ (%) | 3.2 (2.2) | 5.4 (3.6) | 4.8 (3.4) | 2.2 (1.7) | 2.8 (1.7) |
| Median area, $\mu m^2$ (%) | 148 | 148 (0) | 143 (-3.4) | 126 (-14.9) | 161 (8.8) |

**Supplementary Table 5**. Estimated values of VLF amplitude, LF amplitude and median area of penetrating arteriole PVS during sleep states obtained from linear mixed effects models. For VLF amplitude and LF amplitude, numbers in brackets show the percentage wise change in area from median area in each sleep cycle state. For the median area, numbers in brackets show percentage wise change in area from wake-before-sleep state.

| **Endfoot tube diameter** | Wake before sleep | NREM | IS | REM | Wake after sleep |
|---|---|---|---|---|---|
| VLF amplitude, µm (%) | 0.19 (1.0) | 0.37 (1.9) | 0.33 (1.7) | 0.21 (1.0) | 0.15 (0.7) |
| LF amplitude, µm (%) | 0.13 (0.7) | 0.24 (1.3) | 0.22 (1.1) | 0.1 (0.5) | 0.13 (0.6) |
| Median diameter, µm (%) | 19 | 19.2 (1.1) | 19.4 (2.1) | 20.5 (7.9) | 19.2 (1.1) |

**Table 6**. Estimated values of VLF amplitude, LF amplitude and diameter of penetrating arteriole endfoot tube during sleep states obtained from linear mixed effects models. For VLF amplitude and LF amplitude, numbers in brackets show the percentage wise change in diameter from median diameter in each sleep cycle state. For median diameter, numbers in brackets show percentage wise change in diameter from wake-before-sleep state.

| **Lumen diameter** | Quiet wakefulness | Whisking | Locomotion |
|---|---|---|---|
| VLF amplitude, µm (%) | 0.5 (4.4) | 0.4 (3.3) | 0.63 (4.7) |
| LF amplitude, µm (%) | 0.37 (3.2) | 0.31 (2.5) | 0.46 (3.5) |
| Median diameter, µm (%) | 11.4 | 12.3 (7.9) | 13.3 (16.7) |

**Supplementary Table 7.** Estimated values of VLF amplitude, LF amplitude and diameter of penetrating arteriole lumen during wake states obtained from linear mixed effects models. For VLF amplitude and LF amplitude, numbers in brackets show the percentage wise change in diameter from median diameter in each wakefulness state. For median diameter, numbers in brackets show percentage wise change in diameter from quiet wakefulness state.

| **PVS total width** | Quiet wakefulness | Whisking | Locomotion |
|---|---|---|---|
| VLF amplitude, µm (%) | 0.31 (4.8) | 0.26 (4.3) | 0.57 (10.4) |
| LF amplitude, µm (%) | 0.23 (3.5) | 0.24 (4.0) | 0.44 (8.0) |
| Median total width, µm (%) | 6.5 | 6 (-7.7) | 5.5 (-15.4) |

**Supplementary Table 8**. Estimated values of VLF amplitude, LF amplitude and total width of penetrating arteriole PVS during wake states obtained from linear mixed effects models. For VLF amplitude and LF amplitude, numbers in brackets show the percentage wise change in total width from median total width in each wakefulness state. For total width, numbers in brackets show percentage wise change in total width from quiet wakefulness state.

| **PVS area** | Quiet wakefulness | Whisking | Locomotion |
|---|---|---|---|
| VLF amplitude, $\mu m^2$ (%) | 5.7 (3.9) | 5.2 (3.7) | 10 (7.4) |
| LF amplitude, $\mu m^2$ (%) | 3.9 (2.6) | 4.9 (3.5) | 10 (7.4) |
| Median area, $\mu m^2$ (%) | 148 | 142 (-4.1) | 136 (-8.1) |

**Supplementary Table 9**. Estimated values of VLF amplitude, LF amplitude and median area of penetrating arteriole PVS during wake states obtained from linear mixed effects models. For VLF amplitude and LF amplitude, numbers in brackets show the percentage wise change in area from median area in each wakefulness state. For the median area, numbers in brackets show percentage wise change in area from quiet wakefulness state.

| **Endfoot tube diameter** | Quiet wakefulness | Whisking | Locomotion |
|---|---|---|---|
| VLF amplitude, µm (%) | 0.13 (0.7) | 0.18 (1.0) | 0.3 (1.6) |
| LF amplitude, µm (%) | 0.13 (0.7) | 0.14 (0.8) | 0.32 (1.7) |
| Median diameter, µm (%) | 18.3 | 18.6 (1.6) | 19 (3.8) |

**Supplementary Table 10**. Estimated values of VLF amplitude, LF amplitude and diameter of penetrating arteriole endfoot tube during wake states obtained from linear mixed effects models. For VLF amplitude and LF amplitude, numbers in brackets show the percentage wise change in diameter from median diameter in each wakefulness state. For median diameter, numbers in brackets show percentage wise change in diameter from quiet wakefulness state.

### Supplementary Videos

**Supplementary Video 1**. Two-photon recording of a penetrating arteriole in layer II/III somatosensory cortex across a sleep cycle. Scale bar 20 µm.
https://drive.google.com/file/d/1MYrvXT3Ip9pOJQ7CGVlXpAMG1o0j60N1/view?usp=sharing
